# Age-dependent roles of cardiac remodeling in sepsis defense and pathogenesis

**DOI:** 10.1101/2023.03.14.532695

**Authors:** Karina K. Sanchez, Justin L. McCarville, Sarah J. Stengel, Jessica M. Snyder, April E. Williams, Janelle S. Ayres

**Author notes:** These authors contributed equally (Karina K. Sanchez, Justin L. McCarville).

## Abstract

Disease tolerance is a defense strategy essential for survival of infections, limiting physiological damage without killing the pathogen. The disease course and pathology a pathogen may cause can change over the lifespan of a host due to the structural and functional physiological changes that accumulate with age. Since successful disease tolerance responses require the host to engage mechanisms that are compatible with the disease course and pathology caused by an infection, we predicted that this defense strategy would change with age. Animals infected with a lethal dose 50 (LD_50_) of a pathogen often display distinct health and sickness trajectories due to differences in disease tolerance, and thus can be used to delineate tolerance mechanisms. Using a polymicrobial sepsis model, we found that despite having the same LD_50_, old and young susceptible mice exhibited distinct disease courses. Young survivors employed a cardioprotective mechanism via FoxO1-mediated regulation of the ubiquitin-proteosome system that was necessary for survival and protection from cardiomegaly. This same mechanism was a driver of sepsis pathogenesis in aged hosts, causing catabolic remodeling of the heart and death. Our findings have implications for the tailoring of therapy to the age of an infected individual and suggest that disease tolerance alleles may exhibit antagonistic pleiotropy.

## Introduction

Hosts have evolved two distinct defensive health strategies to protect against infections that can be classified by their effects on pathogen fitness^1,2^. Antagonistic defense strategies protect the host by having a negative impact on pathogen fitness. This includes resistance mechanisms mediated by the immune system that involves pathogen detection and elimination. Physiological defense strategies mediate host adaptation to the infection, yielding an apparent cooperation between the host and pathogen^3,4^. This includes disease tolerance mechanisms that enable the host to withstand the presence of a pathogen by limiting physiological damage that occurs during an infection without killing the pathogen^1,2^. The traditional paradigm for infectious disease research is that pathogen elimination necessarily restores health and survival of a host, and thus the field has primarily focused on understanding mechanisms of immune resistance. However, we intuitively know that survival of infections is not so simple, and is also dependent on limiting tissue damage and sustaining physiological function. Currently our primary methods for achieving this are with supportive care, however a better understanding of the mechanisms that promote disease tolerance may result in new therapeutic options for the treatment of infectious diseases that can complement our current antimicrobial based strategies.

Organismal aging complicates host defense against infection. A decline in the ability to kill pathogens due to a decline in immune function and resistance mechanisms is one of the most well recognized consequences of aging^5,6^. However, we have relatively no appreciation of how aging affects the ability of a host to mount an effective disease tolerance response and adapt to an infection. The distinct specificities of resistance and disease tolerance highlights the special challenges that aging imposes on a host’s ability to cooperate with a pathogen. The specificity of resistance is defined by the pathogen, which is independent of host age. By contrast, the specificity of disease tolerance is defined by the physiological perturbations or pathology that occur in response to the infection, which can theoretically change with age^3,7^. This is because infectious disease pathogenesis is dependent on both the type of pathogen and the host response to the pathogen. As structural and functional changes accumulate with age, how the host responds to an infection can change, which will affect the manner in which pathology occurs and the type of pathology that occurs. This can result in two hosts of different ages exhibiting distinct disease courses and pathologies despite being infected with the same pathogen. This leads to the prediction that in addition to mechanisms of infection pathogenesis being distinct, the disease tolerance mechanisms that are necessary to facilitate physiological adaptation to infection and to protect against pathology will be different in an aged host compared to a young host for the same infection challenge.

Here, we describe our approach to define the consequences of aging for disease tolerance and host-pathogen cooperation. Leveraging the phenomenon of lethal dose 50 (LD_50_) and a polymicrobial sepsis model, we found that while the polymicrobial sepsis LD_50_ dose was the same for both old and young mice, the young mice that succumb to the challenge exhibited a disease course that was distinct from that exhibited by old dying infected mice. In young mice, survivors of the LD_50_ employed a cardioprotective disease tolerance mechanism, mediated by FoxO1-dependent activation of muscle-specific E3 ubiquitin ligases regulating cardiac remodeling and preventing sepsis-induced cardiomegaly and death. By contrast, this same mechanism contributed to sepsis pathogenesis and death in aged hosts by driving cardiac muscle wasting. Our study provides a conceptual advancement for our understanding of the relationship between aging, infectious disease pathogenesis and host adaptation to infection, and reveals a novel mechanism of age-dependent sepsis pathogenesis and disease tolerance. Our findings have implications for the tailoring of therapy to the age of an infected individual and suggest that disease tolerance alleles may exhibit antagonistic pleiotropy.

## Results

### An LD_50_ approach for the investigation of the relationship between disease tolerance and aging

LD_50_ describes the dose of a pathogen that will kill 50% of a genetically identical host population. We previously demonstrated that this approach can be used to elucidate mechanisms of physiological defenses^8^. We hypothesized that this phenomenon would be useful for understanding how aging influences the ability of a host to cooperate with a pathogen. To test our hypothesis, we used a polymicrobial sepsis model consisting of a 1:1 mixture of the Gram negative bacterium *Escherichia coli* and the Gram positive bacterium *Staphylococcus aureus* in 12-week and 75-week old mice. We utilized 12-week old mice because they fall into the mature adult category (20-30 year old humans), and 75-week old mice meet the definition of old (56-69 year old humans)^9^. Mice in these two age groups differ significantly in all homeostatic variables and parameters measured (**Supplemental Figure 2**), demonstrating that the old mice have undergone homeostatic reprogramming. We performed dose titration experiments in young and old mice to determine the LD_50_ dose of polymicrobial sepsis for each age group. In young mice, we found the LD_50_ dose to be ~1×10^8^ total CFU, with 50% of mice succumbing to the infection within the first ~14-hours post infection (**Figure 1A**). Surprisingly, in old mice, we found the LD_50_ dose of polymicrobial sepsis to be the same, with 50% of old mice succumbing to the infection within the first ~26-hours post infection (**Figure 1B**). This suggests that in the context of this infection and within the age range examined, lethality does not increase with age.

**Figure 1.**
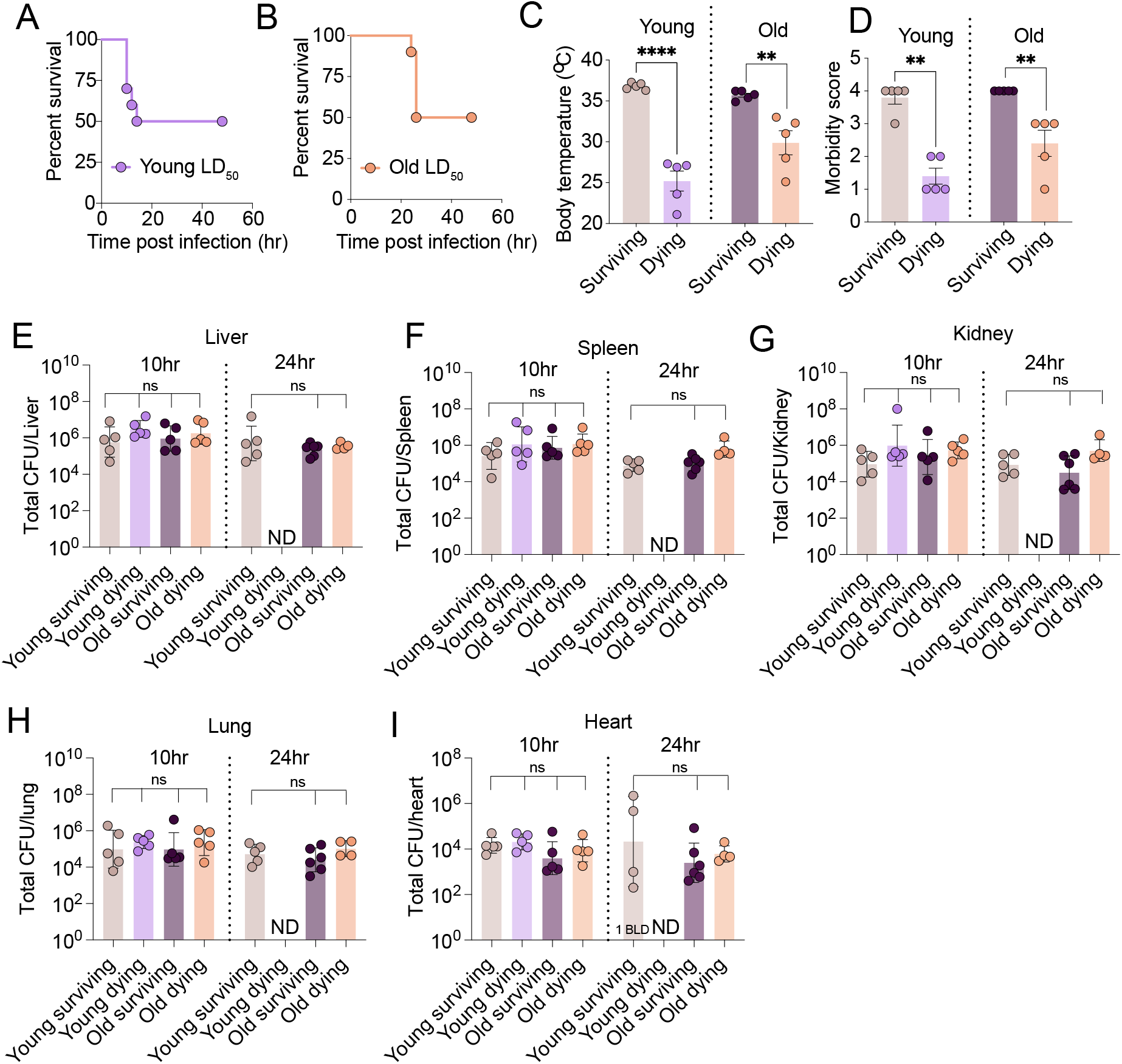
An LD_50_ approach for the investigation of disease tolerance. (A) 12-week old mice were i.p. infected with the LD_50_ dose of polymicrobial sepsis and survival was monitored. n = 10. (B) 75-week old mice were i.p. infected with the LD_50_ dose of polymicrobial sepsis and survival was monitored. n = 10. (C-D) Young and old mice were challenged with LD_50_ dose of polymicrobial sepsis. Temperature and morbidity were quantified every two hours post-infection. (C) Temperature and (D) Morbidity at 10 hours post-infection for young mice and 24 hours post-infection for old mice. Time points represent time at which dying mice from each age group reached clinical endpoint. n = 5 mice per condition. \*\**p* < 0.01, \*\*\*\**p* < 0.0001. (E-I) Young and old mice were challenged with the LD_50_ dose of polymicrobial sepsis and total pathogen burdens (colony forming units, CFU) were determined in (E) Liver; (F) Spleen; (G) Kidney; (H) Lung and (I) Heart at 10 and 24 hours post-infection. ND indicates not determined due to all young dying mice succumbing to the infection prior to 24 hours post-infection. For heart, one mouse below the limit of detection and indicated by “1 BLD”. n = 4-6 mice per group. unpaired t-test, Mann-Whitney test, One-way ANOVA, Kurskal-Wallis test with post Benjamini, Krieger and Yekutieli post-test for pairwise comparisons. Error bars indicate +/- SEM. For CFU plots showing geometric mean +/- SD. Dying mice data from (A-B) also shown in Figure 2A.

While parameters such as lean and fat body mass prior to the infection were not useful for predicting the health trajectories of LD_50_-challenged animals (**Supplemental Figure 3**) for both young and old mice, those that succumb to the LD_50_-challenge developed quantifiable clinical signs of disease that enabled the determination of the fates of each LD_50_-challenged animal prior to any deaths occurring. This included severe hypothermia, as well as severe morbidity including ruffled fur and lack of voluntary movement in the presence or absence of stimuli (**Figure 1C** and **D**). Survivors of the LD_50_ for both age groups were clinically healthy at the time the age-matched LD_50_-challenged dying mice succumb to the infection (**Figure 1C** and **D**). Interestingly, housing mice at thermal neutrality was not sufficient to protect mice from polysepsis induced lethality in old or young mice (**Supplemental Figure 4**). To determine the relationship between infection outcome and pathogen burdens, we challenged young and old mice with the LD_50_ dose of polymicrobial sepsis, assigned them clinical scores every two-hours post-infection based on temperature and morbidity, and assessed pathogen burdens when the dying animals reached maximal morbidity (~10hrs for young mice and ~24hrs for old mice, **Figure 1C-D**). For old mice, the fates for each LD_50_-challenged mouse could be determined as early as ten hours post-infection due to differences in clinical scores, enabling the quantification of pathogen burdens at the early stage of infection. In young mice, we found comparable levels of total bacteria, *E. coli* and *S. aureus* in all organs assayed from the survivors and dying hosts (**Figure 1E-I, Supplemental Figure 5-6**). By our definitions, the differences in infection outcome for young LD_50_-challenged mice are due to differences in the ability to adapt to the infection via disease tolerance mechanisms, rather than differences in the ability to kill the pathogen with resistance defenses^1–3^.

Finally, from our CFU analyses, we determined that old and young survivors of the LD_50_ did not differ in their ability to control the infection levels via resistance defenses, nor did we detect differences in resistance defenses between those that succumb to the LD_50_ as indicated by comparable pathogen burdens between the two groups (**Figure 1E-I, Supplemental Figure 5-6**). This indicates that age related changes to the immune response did not affect the ability of host to control pathogen burdens in this infection model. Taken together, our data demonstrate that because there are no differences in the infectious inoculum between age groups, that there are no apparent age related differences in resistance defenses between the two different age groups that are necessary for controlling this infection, and because the differences in the outcome groups for both old and young mice are due to differences in disease tolerance, our LD_50_ polymicrobial sepsis model is an ideal system to elucidate the principles and mechanisms of how aging influences disease tolerance defenses.

### Aging changes the disease course of LD_50_-challenged mice

Despite having the same LD_50_ dose of polymicrobial sepsis, the disease courses were distinct for old and young mice that died from the LD_50_-challenge. Old dying mice exhibited slower death kinetics compared to young mice (**Figure 2A**). Consistent with this, while old and young dying mice exhibited comparable severity in their hypothermic responses and morbidity (**Supplemental Figure 7A-B**), yielding comparable minimal health scores during their infection course (**Supplemental Figure 7C**), the kinetics of disease progression were distinct, with young dying mice exhibiting faster kinetics than old dying mice (**Figure 2B-C, Supplemental Figure 7D-I**). The delay in clinical manifestations between the age groups was independent of pathogen burdens (**Figure 1E-I Supplemental Figure 5-6**). This suggests that the differences in the disease courses were not due to differences in resistance defenses and the ability to control the infection. Instead, our data suggest that the effects of age-related changes to host physiology led to differences in sepsis pathogenesis and progression. To determine this, we asked if there were any differences in the type or extent of organ damage. While both young and old dying mice exhibited elevated levels of markers for renal and hepatic damage (**Supplemental Figure 8A-C**), we observed age-related differences in sepsis-induced cardiac remodeling phenotypes. Young dying mice exhibited cardiomegaly characterized by a significant increase in heart weight, changes in geometric shape, and histological evidence of edema, changes in cardiomyocyte appearance and hypereosinophilia with increased cardiomyocyte size and leukocyte infiltration (**Figure 2D-J** and **Supplemental Figure 8D**). Old dying mice exhibited severe cardiac atrophy with a significant loss of heart weight and changes in geometric shape (**Figure 2D-F** and **Supplemental Figure 8D**). The rapid kinetics of cardiac remodeling we observed in our mouse models is comparable to what has been reported in humans^10,11^. Histopathological analysis revealed that hearts from old dying mice exhibited similar, although less pronounced, histological changes in cardiomyocyte appearance and leukocyte infiltration indicating that similar acute microscopic changes are associated with different cardiac remodeling at the macroscopic level in young and old dying septic hosts, perhaps consistent with a limited spectrum of histologic changes in the acute time frame of the disease (**Figure 2D-J** and **Supplemental Figure 8D**). Furthermore, RNAseq analysis revealed distinct molecular signatures indicative of cardiac remodeling. Young dying mice had enrichment of genes involved in apoptosis, negative regulation of protein catabolic process, response to muscle stretch, regulation of cell shape and cardiac muscle tissue morphogenesis which is consistent with cardiac remodeling and cardiomegaly (**Figure 2K** and **Supplemental Figure 8E-G**). The cardiac transcriptome of old dying mice had enrichment of genes involved in ubiquitination, autophagic cell death, negative regulation of cell growth, cardiac muscle development and cell proliferation, which is consistent with their cardiac atrophic remodeling phenotype (**Figure 2K** and **Supplemental Figure 8E-G**). Gene ontology analysis also revealed that old and young dying mice had distinct gene depletion profiles (**Supplemental Figure 8G**). Young, but not old dying, mice exhibited elevated levels of Troponin I, a marker of cardiac damage (**Supplemental Figure 8H**).

**Figure 2.**
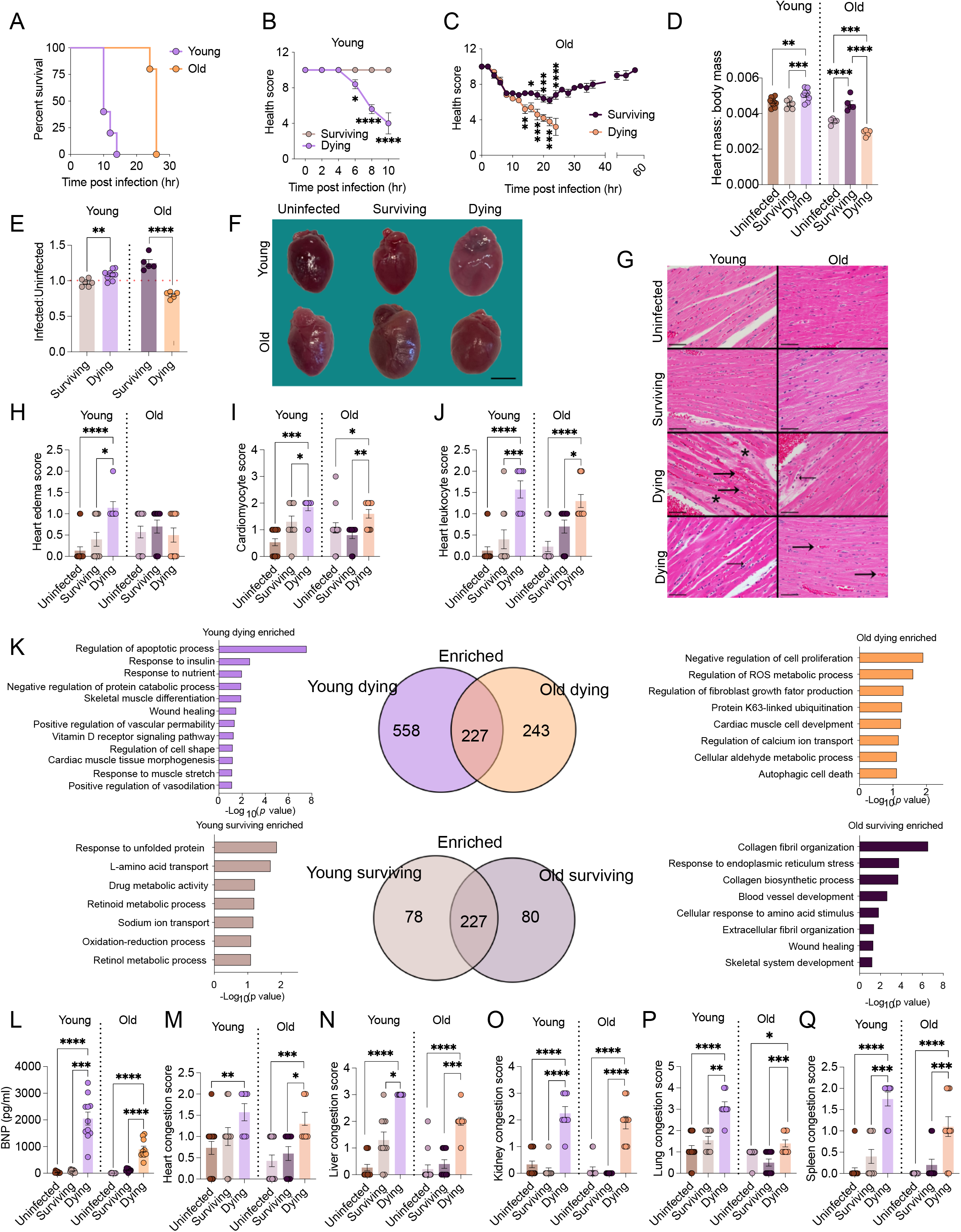
Age dependent health and disease trajectories exhibited by polymicrobial sepsis LD_50_ challenged mice. (A) Death kinetics of old and young mice from Figure 1A-B that succumb to the LD_50_ polymicrobial sepsis challenge. n = 5 mice per condition, *p* = 0.0015. (B-C) 12-week and 75-week old mice were challenged with the LD_50_ dose of polymicrobial sepsis. Temperature and morbidity were measured every two hours to calculate health scores. Health scores over time are shown to determine health and disease trajectories of surviving and dying mice. (B) Trajectories of young mice. (C) Trajectories of old mice. n = 5 mice per group. \*\*\*\**p*<0.0001, \*\*\**p*<0.005, \*\**p*<0.01, \**p*<0.05. Plots with individual data points are shown in Supplemental Figure 7F and I. (D-J) 12-week and 75-week old mice were challenged with the LD_50_ dose of polymicrobial sepsis. Hearts were harvested for macroscopic and microscopic analyses. (D) Heart weights normalized to body weight. (E) Values of infected mice from (D) normalized to average of uninfected values from (D). n = 5-10 mice per condition. \*\*\*\**p*<0.0001, \*\*\**p*<0.005, \*\**p*<0.01, \**p*<0.05. (F) Representative images of hearts. Original images in Supplemental Figure 8D. Scale bar indicates 3mm. (G) Representative images of heart at the level of the interventricular septum in young mice (left column) and old mice (right column). From top to bottom: uninfected mice (top row); infected surviving mice (second row); and infected dying mice (bottom two rows). Compared to uninfected mice and infected surviving mice, infected dying mice have more prominent myocardial vessels with increased red blood cells (congestion, large arrows) and leukocytes (small arrows), mildly increased hypereosinophilic cardiomyocytes and edema (asterisks). Scale bar indicates 50 microns. (H) Heart edema score, (I) Cardiomyocyte score, (J) Heart leukocytes score from histopathology analysis. 7-15 mice per condition. Two experiments combined. \*\*\*\**p*<0.0001, \*\*\**p*<0.005, \*\**p*<0.01, \**p*<0.05. (K) Young and old mice were challenged with the LD_50_ dose of polymicrobial sepsis. When dying mice for each age group reached minimal health (clinical endpoint), hearts were harvested from age-matched infected surviving and uninfected control mice and RNAseq was done (10 hrs post-infection for young mice and 24 hrs for old mice). Venn diagrams of genes that were enriched in young dying hearts compared to both young surviving and young uninfected mice, vs old dying hearts compared to both old surviving and old uninfected mice, and young surviving hearts compared to both young dying and young uninfected mice vs old surviving hearts compared to both old dying and old uninfected mice. Gene ontology analysis was done on the distinct gene lists for each group as shown in the bar graphs. (L) Serum BNP levels of LD_50_-challenged and uninfected age matched mice. n = 9-11 mice per condition. \*\*\*\**p*<0.0001, \*\*\**p*<0.005. (M-Q) Histopathology analysis of organs from LD_50_-challenged and uninfected age matched mice. (M) Heart congestion, (N) Liver congestion, (O) Kidney congestion, (P) Lung congestion, (Q) Spleen congestion. n = 7-15 mice per condition. Two experiments combined. \*\*\*\**p*<0.0001, \*\*\**p*<0.005, \*\**p*<0.01, \**p*<0.05. Error bars indicate +/-SEM. For pairwise comparisons, One-way ANOVA or Kruskal-Wallis test with Two-stage linear step-up procedure of Benjamin, Krieger and Yekutieli, or Two-way ANOVA.

However, dying mice of both age groups exhibited elevated levels of BNP and Galectin 3, which are markers of cardiac failure (**Figure 2L** and **Supplemental Figure 8I**). Finally, both old and young dying mice exhibited congestion of vital organs, which can be indicative of cardiac failure (**Figure 2M-Q** and **Supplemental Figure 8J**). Taken together, our data show that despite being infected with the same type and dose of pathogen, and achieving comparable infection levels, aging causes infected dying animals to exhibit distinct disease courses prior to death that involves opposing cardiac remodeling phenotypes.

### Aging changes the health trajectories of LD_50_-challenged mice

The distinct routes to death suggested that what is important for survival of the LD_50_ would be different for old and young mice. We hypothesized that this would be reflected by the health trajectories of the survivors of the LD_50_. Indeed, LD_50_-challenged young survivors were able to maintain their body temperature and did not exhibit clinical signs of disease over the course of the infection, which is indicative of an endurance phenotype (**Figure 2B** and **Supplemental Figure 7A-F**). By contrast, old LD_50_-challenged survivors exhibited a resilience health trajectory, with a sickness phase characterized by morbidity and hypothermia that plateaued at ~6-8hrs post-infection, and a return to baseline health between 24-48hrs post-infection (**Figure 2C** and **Supplemental Figure 7A-C, G-I)**. The differences in the health trajectories were independent of pathogen burdens, indicating that the ability to control sickness in young survivors compared to the old survivors was not due to differences in resistances defenses (**Figure 1E-I Supplemental Figure 5-6**). Furthermore, our CFU analysis demonstrated that the bifurcation of the health trajectories of old surviving and dying mice at ~10hrs postinfection was not due to differences in resistance defenses and the ability to control infection levels **Figure 1E-I, Supplemental Figure 5-6**). Old surviving mice were protected from both kidney and liver damage, while young survivors were protected from kidney damage, and also mildly protected from liver damage (**Supplemental Figure 8A-C**). Young survivors were protected from sepsis induced cardiomegaly, edema, changes in cardiomyocyte appearance, and leukocyte infiltration and instead exhibited a trend toward mild cardiac atrophy compared to uninfected young mice (**Figure 2D-J** and **Supplemental 9D**). Old survivors exhibited cardiomegaly without histological changes in edema and cardiomyocyte appearance, or a significant increase in leukocyte infiltration (**Figure 2D-J** and **Supplemental Figure 8D**). Our RNAseq analysis revealed that the cardiac transcriptomes of the old and young survivors of the LD_50_ were distinct. Furthermore, they were different from age-matched LD_50_-infected dying and uninfected control mice (**Supplemental Figure 8E**). This demonstrates that there are distinct cardiac transcriptomes for young and old infected mice that correlate with survival, and that both old and young survivors do not resemble age matched uninfected healthy controls. Gene ontology analysis showed that young survivors had an enrichment of genes involved in the unfolded protein response, amino acid transport, retinoid/retinol process and sodium ion processes (**Figure 2K and Supplemental Figure 8G**). By contrast the old survivors of the LD_50_ had enrichment of genes involved in the regulation of collagen fibril organization and biosynthetic processes, wound healing, cellular response to amino acid and response to endoplasmic reticulum stress (**Figure 2K and Supplemental Figure 8G**). Finally, both old and young survivors were protected from cardiac damage and failure (**Figure 2L-Q** and **Supplemental Figure 8H-J**,). Thus infected surviving animals exhibit age-specific and distinct health trajectories despite being infected with the same dose and pathogen. Finally, our results suggest that cardiac remodeling has opposing costs for infection defense in an age dependent manner. In young mice, protection from cardiomegaly is adaptive. In old mice, allowing cardiomegaly to occur to prevent cardiac muscle wasting is beneficial for host defense.

### FoxO1 promotes a cardioprotective disease tolerance mechanism in young hosts

Young LD_50_-challenged dying animals exhibited cardiomegaly, cardiac damage and failure, but old LD_50_-challenged dying mice exhibited cardiac failure associated with cardiac muscle wasting (**Figure 2D-Q, Supplemental Figure 8D-J**). We hypothesized that a cardioprotective disease tolerance mechanism was operating in the young survivors of the LD_50_ that was not necessary for resilience and survival in the old LD_50_-challenged mice, and that this would be reflected by the cardiac transcriptomes. Disease tolerance mechanisms are a component of the inducible host defense response to an infection and previous studies have demonstrated that relatively minor changes in gene expression can reveal factors important for the regulation of disease tolerance^3,12^. From our RNAseq analysis, we revealed 316 genes that were significantly upregulated in the young LD_50_-challenged survivor hearts compared to the other five animal groups (**Figure 3A-B**). KEGG pathway analysis of this inducible expression signature, revealed the enrichment of various metabolic, cell cycle/survival, muscle and hormone signaling processes, with FoxO signaling pathways being the most significantly enriched pathway (**Figure 3C**). While cardiac Foxo1 expression and activity were not different between old and young animals under uninfected conditions (**Supplemental Figure 9A-C**), we found cardiac FoxO1 was differentially regulated in old and young hosts when challenged with sepsis and in a fate specific manner. Cardiac *Foxo1* expression was significantly induced in young LD_50_-challenged survivors compared to uninfected young mice, young LD_50_-challenged dying mice, as well as old uninfected, dying and surviving mice (**Figure 3D** and **Supplemental Figure 9D**). Protein levels of FoxO1 were also elevated without an increase in the levels of phosphorylated FoxO1, indicating there was increased cardiac FoxO1 activity in young surviving septic mice compared to all other conditions (**Supplemental Figure 9E**). Examination of other vital organs revealed the association between *Foxo1* induction and survival in young mice to be specific to the heart (**Supplemental Figure 9D**). In old mice, survival of sepsis did not correlate with increased expression or activation of FoxO1 in the heart or any other vital organ (**Figure 3D, Supplemental Figure 9D-E).**

**Figure 3.**
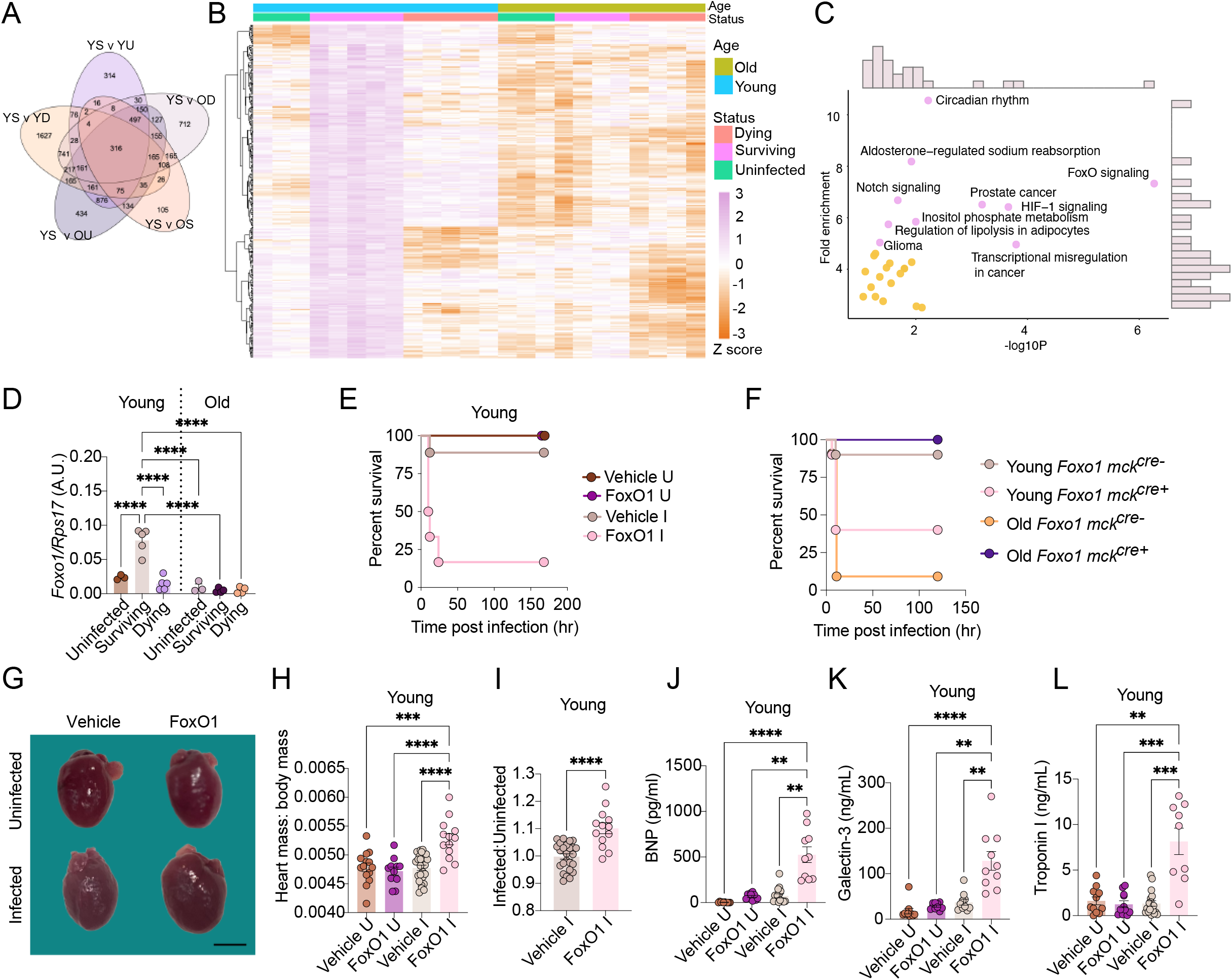
FoxO1 promotes a cardioprotective disease tolerance mechanism in young hosts. (A-C) The cardiac transcriptomes of young and old uninfected and LD_50_ challenged mice were determined as described in Figure 2K. (A) Venn diagram of the genes induced in young surviving mice compared to all other five conditions are shown; YS – young surviving YU – young uninfected, YD – young dying, OS – old surviving, OU – Old uninfected, OD – old dying. (B) Heat map of the overlap of genes for all five comparisons in (A); (C) KEGG pathway enrichment analysis of the 316 genes that were specifically induced in the young surviving mice compared to the other five conditions shown in (A-B) showing 10 most significant groups. (D) Cardiac gene expression of *Foxo1* in uninfected and LD_50_ challenged young and old mice when dying mice in each age group reached maximal morbidity (10hrs for young and 24hrs for old). n = 3-5 mice per condition. \*\*\*\**p* < 0.0001. Data are also shown in Supplemental Figure 9F. (E) Young mice were infected with a low dose of polymicrobial sepsis and treated with a FoxO1 inhibitor or vehicle and survival was measured. *p* value is comparing vehicle I to FoxO1 I; n = 5-9 mice per condition. *p* = 0.0003. (F) Young and old *Foxo1 mck^cre-^* and *mck°^re+^* mice were infected with polymicrobial sepsis and survival was monitored. n = 9-11 mice per condition. *p* = 0.0212 for young and *p* = 0.0001 for old. (G) Representative heart images from young uninfected and vehicle infected surviving and FoxO1 inhibitor treated dying mice at 10hrs post-infection. Original images are shown in Supplemental Figure 11A. Scale bar – 3mm. (H) Heart weights normalized to body weights from weight-matched young uninfected and infected mice treated with a FoxO1 inhibitor or vehicle at 10hrs post-infection. n = 11-25 mice per condition. \*\*\**p* < 0.0005, \*\*\*\**p* < 0.0001. (I) Values of infected mice from (H) normalized to the average of uninfected values from (H). n = 13-25 mice per condition. \*\*\*\**p* < 0.00001. (J) Serum BNP levels of young uninfected, vehicle infected surviving and FoxO1 inhibitor treated dying mice at 10hrs postinfection. n = 10-15 mice per condition. \*\**p* < 0.005, \*\*\**p* < 0.0005 *\*\*\*\*p* < 0.0001. (K) Serum Galectin-3 levels of young uninfected, vehicle infected surviving and FoxO1 inhibitor treated dying mice at 10hrs post-infection. n = 10 mice per condition. \*\**p* < 0.01, \*\*\*\**p* < 0.0001. (L) Serum Troponin I levels of young uninfected, vehicle infected surviving and FoxO1 inhibitor treated dying mice at 10hrs post-infection. n = 9-18 mice per condition. \*\**p* < 0.005, \*\*\**p* < 0.0005. Vehicle U – vehicle uninfected, FoxO1 U – FoxO1 inhibitor uninfected, Vehicle I – vehicle infected, FoxO I – FoxO1 inhibitor infected. Error bars +/- SEM. For pairwise comparisons, t-test, One way ANOVA, Kurskal Wallis with Two-stage linear step-up procedure of Benjamini, Krieger and Yekutieli. For survival, Log-rank analysis.

To test the importance of FoxO1 for infection defense, we infected young mice with a low dose of polymicrobial sepsis (LD_10_) and treated with a cell permeable inhibitor of FoxO1 that blocks the transcriptional activity of FoxO1^13^. Inhibition of FoxO1 activity rendered young mice significantly more susceptible to infection-induced sickness and lethality (**Figure 3E**, **Supplemental Figure 10A-C**). This increase in susceptibility was specific to the infected state, as uninfected mice treated with the FoxO1 inhibitor survived and showed no changes in their health (**Figure 3E**, **Supplemental Figure 10A-C**). Furthermore, genetic deletion of *Foxo1* from cardiac muscle rendered young animals highly susceptible to a low dose of polymicrobial sepsis (**Figure 3F**, **Supplemental Figure 10D-H**). We found no differences in *S. aureus, E. coli* or total pathogen burdens in any organs examined in young infected mice treated with the FoxO1 inhibitor or vehicle (**Supplemental Figure 10I-K**). This demonstrates that consistent with our LD_50_ analysis, FoxO1 promotes health and survival of infection in young mice by enabling the host to adapt to the infection rather than by promoting pathogen killing.

Sepsis pathogenesis, and the resulting organ damage and death, is caused by a dysregulated host response to infection^14^. FoxO1 can promote host adaptation to the polymicrobial sepsis infection by changing organ susceptibility to damage via a disease tolerance mechanism^3^. An alternative hypothesis is that FoxO1 activity could be dampening the inflammatory response that causes organ damage and death. We found no differences in the circulating cytokine/chemokine profile between vehicle mice and FoxO1 inhibitor treated mice (**Supplemental Figure 10L-M**), suggesting that FoxO1 mediated protection is largely independent of the magnitude of the inflammatory response. Instead, our data demonstrate that FoxO1 enables the host to adapt to the infection by limiting organ susceptibility to damage via a disease tolerance mechanism. To begin to understand how FoxO1 activity in young mice promotes disease tolerance in response to polymicrobial sepsis infection, we asked if inhibition of FoxO1 activity rendered mice more susceptible to organ damage. We found that the inhibition of FoxO1 activity in young mice led to cardiomegaly characterized by a change in geometrical shape, increased heart weight, and histological changes in cardiomyocyte appearance in the myocardium, edema and leukocyte infiltration, as well as signs of heart damage and failure (**Figure 3G-L, Supplemental Figure 11A-K**). Young FoxO1 inhibitor treated septic mice were not more susceptible to infection induced liver damage, but were more susceptible to kidney damage, however this appears to be independent of the infection as uninfected FoxO1 inhibitor treated mice also exhibited signs of renal damage (**Supplemental Figure 11L-N**).

### FoxO1 regulation of Trim63 protects from sepsis-induced cardiomegaly and death in young hosts

Cardiac remodeling involves molecular, cellular and interstitial changes in the heart that are dependent in part on transcriptional activity that occurs in response to insult^15^. We examined the cardiac expression profile of FoxO1 target genes in young LD_50_-challenged mice and age-matched uninfected controls. We revealed four classes of genes (**Figure 4A**). Class 1 represents genes that are induced in healthy mice independent of infection status. Class 2 represents genes that are induced specifically in healthy-infected mice. Class 3 represents genes that are induced in morbid-infected state. Class 4 represents genes that are induced in infected mice regardless of health status. Within Class 2 were the atrogenes *Trim63* and *Fbxo32*, which are muscle specific transcriptionally regulated E3 ubiquitin ligases that are critical regulators of protein degradation and play an important role in muscle remodeling^16^ (**Figure 4A** and **Supplemental 12A-B)**. We found cardiac *Trim63* and *Fbxo32* to be differentially regulated in old and young hosts when challenged with sepsis and in a fate specific manner. Cardiac *Trim63* and *Fbxo32* expression were significantly induced in the hearts of young LD_50_-challenged survivors compared to uninfected young mice and young LD_50_-challenged dying mice (**Figure 4B-C** and **Supplemental Figure 12C-D**). In old mice, there was no correlation with sepsis outcome and cardiac *Trim63* and *Fbxo32* expression in LD_50_-challenged mice (**Figure 4B-C, Supplemental Figure 12C-D).** Treatment of young infected mice with a FoxO1 inhibitor prevented the induction of both cardiac *Trim63* and *Fbxo32* expression compared to infected vehicle treated controls, demonstrating that FoxO1 activity is necessary for sepsis-induced expression of these enzymes in the heart of young mice (**Figure 4D-E**). We generated mice deficient for *Trim63* to test its importance for infection defense. Young *Trim63* deficient septic mice were highly susceptible to morbidity and mortality when challenged with a low dose of polymicrobial sepsis (**Figure 4F, Supplemental Figure 12E-G**). Consistent with our LD_50_ and FoxO1 models, the increased susceptibility of young *Trim63* deficient mice was associated with the development of cardiomegaly characterized by macroscopic and microscopic changes including geometric shape, weight and edema, as well as signs of cardiac failure (**Figure 4G-L, Supplemental Figure 12H-L, 13A-F**). Young wild type infected mice exhibited significantly lower heart weights, with animals losing about ~20% of their heart mass, demonstrating that cardiac remodeling associated with cardiomegaly is more costly to host health in young septic mice than atrophic cardiac remodeling (**Figure 4G-I** and **Supplemental Figure 12H**). As Trim63 is a muscle specific E3 ubiquitin ligase (**Supplemental Figure 12A**)^16^, we also examined how Trim63 in skeletal muscle was affected by our sepsis model. In contrast to the heart, *Trim63* expression in skeletal muscle did not correlate with survival in young septic mice and was not regulated by FoxO1 during infection (**Supplemental Figure 13G-J**). Consistent with this, Trim63 did not render young septic mice more susceptible to skeletal muscle wasting and skeletal muscle wasting did not correlate with fate in LD_50_-challenged young mice (**Supplemental Figure 13K-R**). Thus sepsis-induced FoxO1 regulation of Trim63 and muscle size is specific to the cardiac muscle. Finally, *Trim63* deficient mice exhibited increased susceptibility to sepsis induced kidney and liver damage (**Supplemental Figure 13S-U**). Taken together, in young mice, FoxO1 promotes disease tolerance via a cardioprotective mechanism mediated by Trim63 that prevents cardiomegaly, cardiac remodeling/damage, morbidity and death.

**Figure 4.**
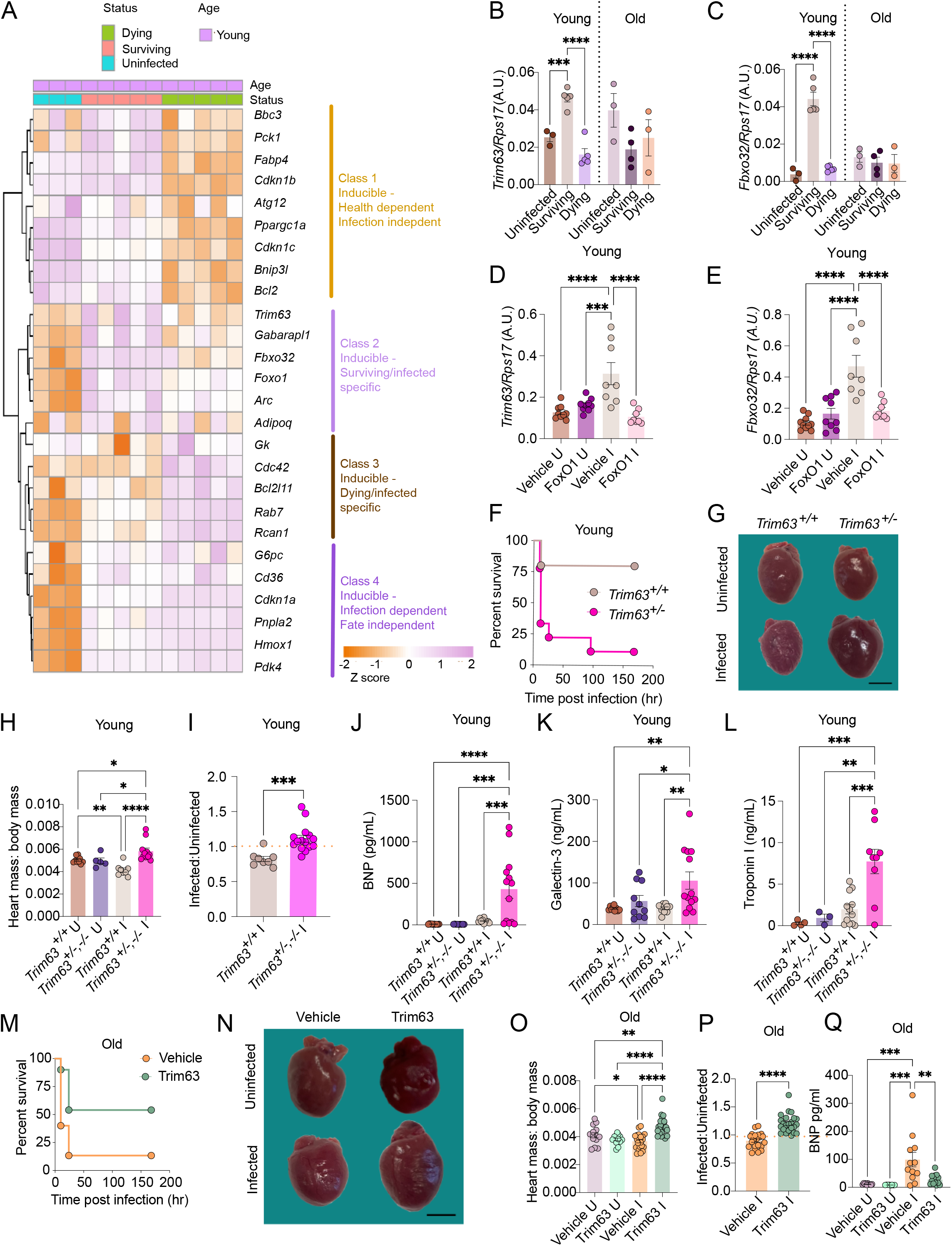
FoxO1 regulation of Trim63 is cardioprotective in young septic mice but pathogenic in old mice. (A) Heat map of FoxO1 targets in hearts from young uninfected and LD_50_-challenged mice from RNAseq analysis presented in Figure 2. n = 3-5 mice per condition. (B-C) 12 and 75 week old mice were infected with the LD_50_ dose of polymicrobial sepsis. Hearts were harvested when dying animals in each age group reached maximal morbidity (10 hrs for young and 24 hrs for old) and expression of (B) *Trim63* and (C) *Fbxo32* were measured. n = 3-5 mice per condition. \*\*\**p* < 0.0005, \*\*\*\**p* < 0.0001. Data are also presented in Supplemental Figure 12C-D. (D-E) Young mice were infected with a low dose (~LD_10_) of polymicrobial sepsis and treated with a FoxO1 inhibitor or vehicle. Hearts were harvested at 10 hr post-infection and expression of (D) *Trim63* and (E) *Fbxo32* were measured. n = 8-10 mice per condition. \*\*\**p* < 0.0005, ****p < 0.0001. (F) Young wild type and *Trim63^+/-^* mice were infected with a low dose of polymicrobial sepsis (~LD25) and survival was determined; n = 5-9 mice per condition. *p* = 0.0186. (G-L) Wild type and *Trim63^+/-,-/-^* mice were infected with a low dose of polymicrobial sepsis (~LD25). Serum and hearts were harvested ~10 hours post-infection. (G) Representative images of hearts. Scale bar = 3mm. Original images are shown in Supplemental Figure 12H. (H) Weights of hearts normalized to body weight. n = 5-11 mice per condition. \**p* < 0.05, **p < 0.01, \*\*\*\**p* < 0.0001. (I) Infected values from (H) normalized to the average of uninfected values from (H). n = 8-15 mice per condition. \*\*\**p* = 0.0007. (J) Serum BNP levels. n = 9-13 mice per condition. \*\*\**p* < 0.0005, \*\*\*\**p* < 0.0001. (K) Serum Galectin-3 levels. n = 9-13 mice per condition. \*\**p* <0.005, \**p* < 0.05. (L) Serum Troponin I levels. n = 3-12 mice per condition. **p < 0.005, \*\*\**p* < 0.0005. (M) Old infected mice were treated with vehicle or a Trim63 inhibitor and survival was monitored. n = 10 mice per condition. *p* = 0.0227. (N-Q) 75 week old mice were infected with polymicrobial sepsis and treated with a vehicle or Trim63 inhibitor. Hearts and serum were harvested at 24hrs post-infection. (N) Images of representative hearts harvested from old infected and uninfected mice treated with vehicle or a Trim63 inhibitor. Scale bar = 3mm. Original images are shown in Supplemental Figure 14F. (O) Weights of hearts normalized to body weight. n = 13-23 mice per condition. \**p* < 0.05, \*\**p* < 0.005, \*\*\*\**p* < 0.0001. (P) Infected values from (O) normalized to the average of uninfected values from (O). n = 21-23 mice per condition. \*\*\*\**p* < 0.0001. (Q) Serum BNP levels. n = 8-13 mice per condition. \*\**p* < 0.005, \*\*\**p* < 0.0005. *Trim^+/+^* U = wild type uninfected; *Trim63^+/-^,^-/-^* U = heterozygous and homozygous Trim63 mutants uninfected; *Trim^+/+^* I = wild type infected; *Trim63^+/--/-^* I = heterozygous and homozygous Trim63 mutants infected. Vehicle U – vehicle treated uninfected, Trim63 U – Trim63 inhibitor treated uninfected, Vehicle I – vehicle treated infected, Trim63 I – Trim63 inhibitor treated infected Error bars indicate +/- SEM. For pairwise comparisons, t- test, One way ANOVA, Kurskal Wallis with Two-stage linear step-up procedure of Benjamini, Krieger and Yekutieli. For survival, Log-rank analysis.

### FoxO1-Trim63 is a driver of sepsis-induced cardiac atrophy and death in old hosts

While we found no association between the expression/activity levels of cardiac FoxO1 and outcome for sepsis infection using our LD_50_ model (**Figure 3D, Supplemental Figure 9F-G)**, because we found catabolic cardiac remodeling old dying septic mice, we wanted to determine if FoxO1-Trim63 contributed to sepsis pathogenesis in aged hosts. Deletion of *Foxo1* from cardiac muscle conferred protection against sepsis induced morbidity and mortality in old mice (**Figure 3F, Supplemental 10D-H**). Additionally, inhibition of FoxO1 activity in wild type mice conferred protection against sepsis induced morbidity and mortality in old mice demonstrating that FoxO1 activity during the infection rather than the accumulation of FoxO1 activity effects over the life of the animal were responsible for the increased susceptibility (**Supplemental Figure 14A-D**). Inhibition of FoxO1 activity in old septic hosts protected from the development of cardiac atrophy and instead resulted in cardiomegaly (**Supplemental Figure 14E-H**). Cardiac *Trim63* expression was induced in both old survivors and dying septic mice at 10hrs post-infection, but returned to baseline levels by 24hrs post-infection (**Supplemental Figure 12C**). Induction at 10hrs was dependent on FoxO1 activity (**Supplemental Figure 14**I). Inhibition of Trim63 activity or genetic deletion of *Trim63* protected old mice from sepsis induced morbidity and mortality (**Figure 4M, Supplemental 14J-P**). Consistent with our LD_50_ model, inhibition of Trim63 activity protected old mice from atrophic cardiac remodeling by promoting cardiomegaly, as well as protection from heart failure (**Figure 4N-Q, Supplemental Figure 14Q-T, 15A-H**). Inhibition of Trim63 did not change the susceptibility of old mice to renal or hepatic damage during infection but did protect from skeletal muscle wasting, however this does not correlate with survival in old septic mice, and was not dependent on FoxO1 (**Supplemental Figure 13O-R, 15I-K, L-Q**). Thus, in contrast to young mice, FoxO1 and Trim63 activity contribute to sepsis pathogenesis in old mice, by driving cardiac muscle wasting, morbidity and death.

## Discussion

Infectious disease pathogenesis is dependent on both the pathogen and the host response to the pathogen. Host factors that dictate whether or not a pathogen will cause disease include: 1) resistance mechanisms that kill the pathogen; 2) the collateral damage of the host response that contributes to pathology; 3) the baseline physiology or vigor of the host when uninfected; and 4) disease tolerance mechanisms that facilitate the ability of the host to adapt to the presence of the pathogen. The aging process will influence each of these host factors. Thus, aging is not only a determinant of whether a host is susceptible to infection, but will also change how a host gets sick and the disease course an individual will take during the infection. While we intuitively realize this, we seldom integrate this way of thinking into our studies of aging and infectious diseases. Instead, we have largely focused on how immune function and the ability of a host to kill a pathogen declines with age. In the present study, using a polymicrobial sepsis model, we took advantage of the curious phenomenon of LD_50_ to ask how aging influences disease tolerance defenses. We found that while the polymicrobial sepsis LD_50_ dose was the same for both old and young mice, the young mice that succumb to the challenge exhibited a disease course that was distinct from that exhibited by old dying challenged mice. Importantly, in our model, in addition to the infectious dose, the pathogen burdens in all target tissues were comparable in the young and old settings, allowing us to conclude that the differences in the disease trajectories were not a function of pathogen burdens. Rather these differences reflect differences in how the host responds to infections due to age-related changes in host physiologies. As sepsis in part is driven by an overexuberant inflammatory response to the infection, our work suggests that the age-related decline in immune function may be sufficient to delay the disease kinetics in aged mice but without compromising resistance to the specific pathogens we tested.

Previous reports of sepsis-induced heart abnormalities in rodents described histologic abnormalities of myofiber arrangement, edema, and inflammatory cell infiltration on HE analysis 6-12 hours following cecal ligation and puncture (CLP)^17,18^, similar to the changes described and scored in both young and old dying mice in the current study. Despite the similar histological changes, we found differences in the cardiac response to injury between aged mice and young dying mice at the macroscopic level. Young dying mice exhibited larger hearts than uninfected or infected surviving controls; whereas, aged dying mice exhibited cardiac muscle wasting and aged surviving mice exhibited larger hearts than uninfected or infected dying controls. The ability of the heart to undergo atrophic remodeling or increase in size is called cardiac plasticity, and enables the heart to change in response to environmental changes. While this can be adaptive, it also can be pathogenic under certain conditions. Indeed, we found that cardiomegaly and atrophic cardiac remodeling have opposing benefits for infection defense in an age dependent manner. In young mice, allowing mild atrophic cardiac remodeling to occur and preventing cardiomegaly is adaptive. In old mice, allowing cardiomegaly is beneficial for host defense. It is unknown precisely how the aging process influences the gross and histological features of sepsis-driven myocardial dysfunction and cardiac hypertrophy, although previous authors have described different histological, ultrastructural and echocardiographic findings in young vs aged mice at 18 hours following CLP^19^. Differences in the histological features in aged mice have also been observed in a genetic model of cardiac hypertrophy. A study by Hao *et al*. determined that osteoprotegerin knock-out (*OPG^-/-^*) mice developed mild cardiac hypertrophy and alterations in the histological presentation of hypertrophy changed throughout the life of these animals. Hypertrophic young *OPG^-/-^* mice had heavier hearts and increased left ventricle wall thickness compared to age-matched wild type mice. However, aged hypertrophic *OPG^-/-^* mice had heavier hearts compared to age matched control mice independent of left ventricle wall thickness, and instead, a greater left ventricle chamber size contributed to the heavier hearts observed^20^. These data and our data suggest the presentation of cardiac hypertrophy can change with age. Future studies should evaluate the relationship between myocyte length and left ventricle chamber size in young and old septic mice to better define the cardiac response to disease in different age groups.

Antagonistic pleiotropy theory describes alleles that are selected for because they have beneficial effects early in life even if these same alleles are detrimental later in life^21^. Resistance mechanisms are often described as exhibiting antagonistic pleiotropy because they are beneficial for fighting infections early in life even if the accumulation of such responses contribute to aging and inflammatory related diseases. Our findings suggest that alleles that contribute to disease tolerance defenses may also exhibit antagonistic pleiotropy. Because the sepsis induced disease courses are distinct in old and young mice, our results show that there can be incompatibilities between disease tolerance mechanisms and disease courses. This would indicate that in some contexts, inappropriate activation of a disease tolerance mechanism may not just be ineffective, but actually may be maladaptive for the host. Opposing roles for disease tolerance mechanisms in host defense have been previously demonstrated in fly studies of young hosts in the context of a diverse array of pathogens, however in these cases there were trade-offs between resistance and tolerance defenses^22,23^. In a mouse study, sickness-induced anorexia was shown to have opposing roles for disease tolerance in bacterial and viral inflammation^24^. As described above, we currently do not know why old and young hosts require different types of cardiac remodeling to defend against sepsis. The cardiovascular system experiences significant changes as an organism ages that will change the way the host responds to an infection and therefore changes the disease course, which can result in the necessity to engage distinct disease tolerance mechanisms. For example, young and elderly septic patients exhibit different hemodynamic characteristics during sepsis^25^. An increase in heart size may be necessary to maintain cardiac output when there is ventricular wall stress or vasodilation. Furthermore, the metabolic demands in old and young septic hosts may be distinct and the protein catabolic response that leads to cardiac atrophy may have opposing roles for host defense in the different age groups.

Previous work has suggested that disease tolerance declines with age^26–28^. In flies, using survival as the readout for health, young flies were only modestly more protected (one day) than old flies when infected with the LD_100_ dose of Flock House virus. There was no difference in viral titers between age groups suggesting that the survival advantage in young flies was due to better disease tolerance^27^. In a mouse model of malaria, 12-month old mice were less resistant to *Plasmodium* compared to 2-month old mice. Using a reaction norm analysis, the authors demonstrated that the old mice were also less tolerant to the parasite only if they used body weight as their readout for health (and not red blood cell count or survival)^28^. With our LD_50_ system, we used survival as our proxy for health and found that the ability to tolerate polymicrobial sepsis infection did not decline with age. We specifically used survival because there are inherent complexities when using clinical phenotypes for which the role of that phenotype in host defense is unknown. Clinical phenotypes such as weight loss, changes in temperature regulation or reduced activity are often viewed as maladaptive and a consequence of the host defense response. However, in some cases these clinical signs of “disease” are a functional component of the host defense response that optimize infection outcome^3^. For example, loss of body weight is typically considered a sign of disease. However, in some context fasted states are necessary for disease tolerance and thus the loss of body fat represents a functional component of the host response to promote survival^23,24^. Thus, conclusions about disease tolerance cannot be drawn based on weight loss alone when the role of the weight loss in promoting host defense is not known. Similarly, in our study, old survivors exhibit mild hypothermia and reduced activity in the early phase of the infection. This may represent a maladaptive consequence of the infection, but it may also represent a component of the defense response that is necessary for protection in the aged host but is not necessary in the young hosts. Until we understand how each of these clinical phenotypes contribute to host defense, they should be used with caution and multiple parameters should be used to draw conclusions about disease tolerance.

We discovered that a FoxO1 dependent cardioprotective mechanism is a component of the inducible host defense response to infection that promotes disease tolerance in an age-dependent manner, by alleviating cardiomegaly cardiac damage and failure. FoxO1 regulates numerous aspects of heart physiology that are critical to defend the heart against many stressors, to prevent pathological conditions including cardiac hypertrophy, via regulation of the ubiquitin ligase system^29^. We demonstrate that protection from cardiac remodeling in young septic hosts is dependent on the E3 ubiquitin ligase, Trim63. Thus, we have revealed a new exciting role for FoxO1 and Trim63 in mediating physiological defense during infection. Whether FoxO1 and Trim63 prevent lethal cardiomegaly in young septic mice by protecting against muscle hypertrophy remains to be determined. Furthermore, Trim63 regulates creatine kinase, a critical regulator of the heart’s energy supply of ATP^30^. Trim63 may therefore promote cardioprotection in young septic mice by regulating metabolism of the heart directly. In old mice, we showed that both FoxO1 and Trim63 contribute to sepsis induced sickness and death and this was associated with cardiac muscle wasting. Increased protein degradation by the ubiquitin-proteosome system in muscle is a common feature of sepsis^31^. This response is best appreciated in the context of skeletal muscle, and is thought to be maladaptive because it can lead to skeletal muscle wasting^32^. Our results suggest that elevated cardiac protein catabolism mediated by the ubiquitin-proteosome system also causes morbidity and mortality in the aged septic host.

Therapeutic strategies for infectious diseases have largely focused on resistance-based strategies and blocking pathogenic responses that lead to physiological damage. The COVID19 pandemic reminds us that our current perspective and strategies for treating infectious diseases are incomplete and highlights the necessity to incorporate the concept of disease tolerance into our studies and treatment of infectious diseases. We often view infectious disease pathogenesis from a single lens and assume the mechanisms that lead to disease are age-independent, leading to the assumption that interventions will be effective irrespective of age. Our findings demonstrate the importance of charting the disease course of different age stages and understanding the health mechanisms that shape those courses. Our findings have important implications for the tailoring of therapy to the age of an infected individual.

## Experimental Procedures

### Mice

For LD_50_ and inhibitor experiments, C57BL/6 female mice from Jackson Labs (000664) were used. Young mice were used at 12-14 weeks of age. Old mice were used at 74-76 weeks of age. *Trim63* mutant mice were generated by Cyagen using CRISPR/Cas mediated genome engineering to delete exons 1-8. The mutant line was maintained as heterozygous and *Trim63^+/-^* and *Trim63^-/-^* were used as *Trim63* KO in our study. The *Trim63^+/+^* generated from the heterozygous breeding scheme were used as wild type controls. *Foxo1 mck^cre^* mice were a generous gift from Dr. Mark Febbraio (Monash University) and were bred in our colony with the following breeding scheme: *Foxo1 mck^cre+^* X *Foxo1 mck^cre-^*. *cre*+ and *cre*-littermates were used for experiments. All experiments were performed in our AAALAC-certified vivarium, with approval from The Salk Institute Animal Care and Use committee (IACUC). In accordance with our IACUC guidelines, in an effort to reduce the number of animals used for our experiments, where possible and appropriate, experiments were done in parallel and shared controls were used. This is indicated in the Figure legends where relevant.

### Bacteria

*E. coli* O21:H+^33^

*S. aureus* (ATCC strain 12600)

## METHOD DETAILS

### Culturing *E. coli O21:H+* and *S. aureus* for mouse infection

*E. coli* O21:H+ was incubated overnight at 37°C on an EMB plate treated with <u>A</u>mpicillin sodium salt (1mg/ml), <u>V</u>ancomycin hydrochloride (.5mg/ml), <u>N</u>eomycin sulfate (1mg/ml), <u>M</u>etronidazole (1mg/ml) antibiotics to grow single colonies. *S. aureus* was incubated overnight on an LB plate at 37°C without antibiotics for colony growth. The next day, a single colony of *E. coli* O21:H+ was inoculated into LB-AVNM media. A single colony of *S. aureus* was inoculated into LB without antibiotics. Both cultures were shaken overnight at 37°C (250 RPM). The following morning, the OD was measured and an inoculum with a 1:1 mixture of the bacteria was prepped with the appropriate amount of both bacteria in sterile 1xPBS that was used directly for mouse infections.

### Mouse infection models

Mice were infected intraperitoneally with the appropriate dose of bacteria and put back into their home cage. Mice were group housed for the experiments. For LD_50_ experiments the dose of total bacteria used was 1×10^8^ CFU. This dose was titrated up or down for low dose and high dose models. Inoculums were serially diluted and plated to confirm the infectious doses. Immediately after infection, food was removed for the first 10-12 hours post-infection to control for any potential variations in the sickness-induced anorexic response. Mice were clinically monitored as described below every two hours post-infection. For some experiments, mice were clinically monitored every two hours for the first 10-12 hrs post-infection and then again at 24hrs. For experiments involving inhibitors, the details are provided below. Mice that reached clinical endpoints were euthanized according to our animal protocol.

### Survival

Mice were clinically monitored as described below every two hours post-infection. For some experiments, mice were clinically monitored every two hours for the first 10-12 hrs post-infection and then again at 24hrs. Mice that had to be euthanized because they reached clinical endpoints during the infection, in addition to those that succumb to the infection, were included in our survival counts.

### Rectal temperature

Rectal temperatures were taken every two hours post infection for the first 10-12 hours, and then every 24 as noted using the Digisense Type J/K/T thermocouple meter. Temperatures are displayed as temperatures over time, temperatures at a defined timepoint or the minimal temperature exhibited by the animals over the course of the infection. This is indicated in the figure legends. Rectal temperatures were also used to calculate health scores to generate health trajectories.

### Grading system for monitoring morbidity

We use the following morbidity scale to quantify the morbidity of mice. Infected mice are clinically assessed using this morbidity scale every two hours post-infection. For some experiments, mice were clinically monitored every two hours for the first 10-12 hrs post-infection and then again at 24hrs. Morbidity scores are displayed as scores over time, scores at a defined timepoint or the maximal morbidity exhibited during the infection (the lower the score, the greater the morbidity). This is indicated in the figure legends. Morbidity scores were also used to calculate health scores to generate health trajectories.

5. Normal. Normal exploratory behavior, rearing on bind limbs, and grooming.
4. Mild. Reduced exploratory behavior, rearing on bind limbs, and grooming. Slower and/or less stead gait, but free ambulation throughout the cage.
3. Moderate. Limited voluntary movement. Slow, unsteady gait got <5 s.
2. Severe. No voluntary movement, but mouse can generate slow, unsteady, gait for > 5 s.
1. Moribund. Mouse does not move away from stimulation by research, but can still right itself.
0. Deceased

### Generating health trajectories

Rectal temperatures were assigned bin scores based on the following strategy. The sum of the temperature bin score and morbidity score (above) for each mouse at each time point was determined to generate health scores. Mice that were deceased or were euthanized because they reached clinical endpoint were assigned a score of 0. Health scores were then plotted against time to generate the health trajectories.

5. 35->38°C
4. 31-34.9°C
3. 28-30.9°C
2. 25-27.9°C
1. 22-24.9°C
0. <22°C

### MRIs

An EchoMRI machine was used for MRI measurements. The MRI produces a total amount of fat (g), total lean muscle (g) and total water (g) per mouse. Both the total fat and lean were normalized to the total body weight (g) of the mouse.

### Heart weights

Weight matched mice were used for heart weight analyses. The body cavity was opened, hearts were removed from the body cavity, blood was drained from the chambers. The heart was then placed on a scale and the weight was recorded. Heart weights were normalized to body weights. For some plots, the value of the heart weight:body weight for infected animals was then normalized to the average heart weight:body weight of the treatment matched uninfected group.

### Heart pictures

For heart pictures, the body cavity was opened, hearts were removed from the body cavity, blood was drained from the chambers and a picture was taken of the heart with a reference ruler using an iPhone SE 2020 or a rose gold iPhone 11 Promax. For presentation purposes, heart images were scaled, cropped and placed on a uniform background using Photopea as follows: Using the reference ruler, each image was scaled to the same size. Hearts were then cropped using a polygonal lasso select tool with one pixel feathering and placed on top of an aquamarine background. A 3mm scale bar is provided in each figure based on the reference ruler in the images. Uncropped images are included in the supplemental figures.

### Histology

Heart, spleen, lungs, liver and kidneys were harvested and fixed in 10% neutral buffered formalin. Lungs were inflated by injecting the lobes with 1xPBS prior to fixation. Then samples were routinely processed, paraffin embedded, sectioned at 4-5 microns, and hematoxylin and eosin stained. Tissues were evaluated by a board certified veterinary pathologist who was blinded to mouse genotype, age, and experimental manipulation, and scored semi-quantitatively for the following parameters: edema (defined as separation of cardiomyocytes by clear to pale eosinophilic material); heart congestion/hemorrhage (defined as blood vessels expanded by erythrocytes in the myocardium and / or extravascular red blood cells); changes to the cardiomyocytes of the myocardium including pallor/cytoplasmic vacuolation, disorganization, and/or hypereosinophilia with or without increased size, loss of nuclear detail, or loss of cross striations; increased leukocytes within the vessels and parenchyma of the heart; spleen congestion (expansion of the red pulp sinuses by erythrocytes); liver congestion/hemorrhage (expanded sinusoids and / or extravascular red blood cells); increased leukocytes within the sinusoids and parenchyma of the liver; congestion and leukocyte infiltration of the lung interstitium; and kidney congestion (expanded vascular spaces in the cortex and medulla). These parameters were scored on a scale of 0-4 with 0 representing normal tissue; 1 represented minimal changes; 2 representing mild changes; 3 representing moderate changes and 4 representing severe changes relative to a score of 1. Representative images were obtained from glass slides using NIS-Elements BR 3.2 64-bit and plated in Adobe Photoshop. Image white balance, lighting, and/or contrast was adjusted using corrections applied to the entire image.

### Quantification of *E. coli* O21:H+ and *S. aureus* in mouse tissues

For quantification of pathogen in tissues, CFUs were quantified. Spleen, kidney, liver, lung, and heart were harvested and homogenized in sterile 1xPBS with 1% Triton X-100 using a BeadMill 24 bench-top bead-based homogenizer (Fisher Scientific). Homogenates were serially diluted and plated on LB agar and EMB-AVNM agar and incubated at 37°C. Colonies were quantified the following day. The limit of detection (LOD) for the liver was 50 CFU and 100 CFU for all other tissues examined. Any sample with values below the LOD (BLD) are indicated on the plots as “X BLD” where “X” indicates the number of mice that had values below the limit of detection for that tissue. Data are plotted as geometric mean +/- SD as indicated in the figure legends.

### Thermal neutrality experiments

The experiment was performed in a temperature controlled housing unit (thermal cabinet), purchased from Columbus instruments (Model #ENC52). Mice were housed in the thermal cabinet set to 30°C, 4 days before infection. At the time of infection, mice were fasted and food was given back 24hrs later. Mouse weight, temperature, and morbidity were measured every two hours during the first day of infection. Mice were tracked for survival on subsequent days. A set of control mice were infected at room temperature with the same dose, fasted for 24hrs, measured every two hours during the first day, and tracked for survival on subsequent days.

### FoxO1 inhibitor infection model

The FoxO1 inhibitor AS1842856 (End Millipore) was injected into mice intraperitoneally at a dose of 40-60mg/kg body weight 36 hours before infection and immediately after infection on the opposite side of bacterial injection. Control mice were injected with vehicle. Mice were infected, clinically monitored and used for downstream analyses as described throughout methods.

### TRIM63 EMBL inhibitor infection model

Old mice were orally gavaged with 30mg/kg body weight of the TRIM63 inhibitor, EMBL (Glixx laboratories) or vehicle as control 12 hours before infection. Mice were then infected with polymicrobial sepsis as described above and two hours post-infection were gavaged with a second dose of EMBL or vehicle. Mice were infected, clinically monitored and used for downstream analyses as described throughout methods.

### Leg muscle measurements

Mice were euthanized and the quadricep, tibialis anterior (TA), extensor digitorum longus (EDL), soleus, and the gastrocnemius were harvested and weighed to determine the mass of each muscle. Muscle weights were then normalized to the body weight and displayed in figures.

### Homeostatic mouse monitoring

For young and old mouse homeostatic mouse data, mice were weighed daily over the course of 10 consecutive days for total body weight tracking. Lean mass, fat mass, free water, and total water were determined using Echo MRI. For food consumption, mice were single caged for 48 hours before food was weighed every 24 hours for two consecutive days at the same time of day. Food consumption over the two days was averaged to report a grams/day average. For rectal temperature, temperature was measured 3 times per day over a consecutive 5-day period at the same time of day using the Digisense Type J/K/T thermocouple meter. The average of the 3 time points per mouse per day were reported. For dissections, mice were euthanized, blood was collected by cardiac puncture, and serum stored at −80°C for future analysis as previously described. Liver, kidney, lung, heart, and spleen were harvested and weighed to determine the mass of each organ.

### Blood pressure

The noninvasive CODA monitor was used to quantify mean blood pressure, systolic blood pressure, and diastolic blood pressure. Mice were placed in a restrainer and warmed on a Far Infrared Warming Platform while recording. The average of 10-15 recordings per mouse per parameter was reported.

### Troponin ELISA

Serum from mice infected with polymicrobial sepsis was harvested by cardiac puncture. Troponin I levels were measured using an ultra-sensitive mouse cardiac troponin-I ELISA kit (Life Diagnostics) according to manufacturer’s protocol. Plates were read on a VERSAmax microplate reader manufactured by Molecular Devices and data analysis was done using SoftMax Pro 5.4.

### *In vivo* metabolic measurements

For the glucose tolerance test (GTT), mice were fasted overnight (12hrs). Tail tips were cut using a razor blade for blood glucose monitoring using the Nova Max Plus glucose monitoring system. Mice were injected intraperitoneally with 2g/kg of glucose (in PBS) based on total body weight. Blood glucose was measured at 0 (before glucose treatment), 15, 30, 45, 60, 90, and 120 minutes post injection. Blood glucose was normalized to percent of time 0, and plotted in Prism where area under the curve was calculated.

### Quantification of whole body metabolic parameters

For quantification of VO2, mice were single caged in metabolic cages within a comprehensive lab animal monitoring system (CLAMS) 24 hours prior to metabolic parameter data collection. Mice were left untouched for approximately 3 days for data collection.

### qPCR

For hearts, livers, spleens, kidneys and lungs, organs were harvested and then frozen at −80°C. They were then homogenized into a tissue powder in liquid nitrogen. The powder was used for subsequent RNA extraction, using the Allprep DNA/RNA Mini Kit (Qiagen) per manufacturer protocol. For leg muscles, the tissues were harvested and then frozen at −80°C. They were then homogenized into a tissue powder in liquid nitrogen. Ground muscle tissue powder was added to cold screw cap tube with bead in liquid nitrogen and then homogenized with 800uL TRIzol LS Reagent (Invitrogen). 160uL of Chloroform (T.C.I.) was added and shaken by hand for 15 seconds and incubated at room temperature for 3 minutes. The Trizol/Chloroform lysate was transferred to 1.5mL tubes and spun in a fume hood centrifuge for 15 minutes at max speed (14,000rpm). The upper aqueous phase was transferred to a new 1.5mL tube and 400uL isopropanol was added and mixed. The samples were held in a −20 freezer for a minimum of 2 hours. The samples were added to an Allprep RNeasy spin column (Qiagen) and spun at max speed for 1 minute, the flow through was discarded. The samples were then processed according to the Allprep RNA (Qiagen) extraction protocol resulting in 60uL of diluted muscle RNA. cDNA was generated using SuperScript IV Reverse Transcriptase (Invitrogen) per manufacturer protocol. qRT-PCR was performed using a QuantStudio 5 Real-Time PCR instrument (Applied Biosystems). Primers sequences are listed in the key resource table. The annealing temperature used was 60°C for *Trim63, Fbxo32*, and *Rps17*. 58°C was used for *Foxo1*.

### BNP and GAL3 quantification

Blood was collected by cardiac puncture and serum was stored at −80°C as described above. For analysis, serum was defrosted and BNP and GAL3 were quantified by a BNP (Cusabio) and GAL3 (Invitrogen) ELISA kit. Samples were diluted at 1:2 for BNP, and at 1:100 to 1:300 for GAL3, and the ELISA was run as specified by both manufacture’s protocols. Plates were read on a VERSAmax microplate reader manufactured by Molecular Devices and data analysis was done using SoftMax Pro 5.4.

### Western Blot Analysis

Heart tissue powder were homogenized in 700uL Tissue Extraction Reagent II supplemented with Protease Inhibitor Cocktail (100:1) using BeadMill 24 bench-top bead-based homogenizer (Fisher Scientific). Lysates were centrifuged at 4°C for 15 min at 27,000rpm and transferred to a new tube. Samples were quantified with a BCA reaction and subjected to western analysis of GAPDH (Cell Signaling), phospho-FoxO1/FoxO3 (Cell Signaling), FoxO1 plus FoxO3 (Cell Signaling). Samples were mixed with 15uL of a 1:10 mixture of 2-Mercaptoethanol to NuPAGE™ LDS Sample Buffer (4X) (Invitrogen), then incubated at 70°C for 10 min and sonicated in a water bath for 10 min. Samples were loaded equally in 7% NuPage 1.0mm x 12 well Tris-Acetate gels with 4°C Tris-Acetate SDS running buffer (50ml 20X Tris Acetate SDS in 950ml DI H2O) for 60 min at 150V. Gels were placed on Trans-Blot^®^ Turbo™ Midi 0.2 μm Nitrocellulose Transfer Packs (Bio-Rad) and inserted into Trans-Blot^®^ Turbo™ Transfer System (Bio-Rad) for 10 min. Membranes were stained with Ponceau Total Protein Stain (Prometheus) to be marked, washed with 1X TBST to de-stain, and placed into blocking solution (5% BSA in 1X TBST) on a shaker platform overnight at 4°C. Blocking solution was removed and primary antibody solutions were added to membranes (1:1000 Ab in Prometheus OneBlock™ Western-CL Blocking Buffer) and placed on shaker platform overnight at 4°C. Primary antibody solutions were removed and membranes were washed on shaker platform for 5 min with 1X TBST, repeated 4 times at RT. Secondary antibody solution was added containing 1:3000 anti-rabbit IgG HRP-linked Ab in blocking buffer (5% BSA in 1X TBST) and membranes were placed on shaker platform for 1hr at RT. Secondary antibody solutions were removed and membranes were washed again on a shaker platform for 5 min with 1X TBST, repeated 4 times at RT. Nitrocellulose blots were developed using a mixture of Femto and Dura chemiluminescent reaction and visualized with a Bio-Rad Gel Doc XR+ machine. For total FoxO1 and pFoxO1 blots, the same lysate was run on two different gels at the same time in the same gel rig. The gels were then transferred to the same membrane. After staining of the membrane with Ponceau, the membranes were cut and destained. The cut membranes were then probed for the relevant protein. pFoxO1 and total FoxO1 blots had their own GAPDH loading control that was run on the same gel.

### Total bicarbonate, potassium, anion gap, cytokine, AST, ALT and BUN quantification by IDEXX

Serums harvested by cardiac puncture were analyzed by IDEXX Bioanalytics.

### RNA-seq and data processing

Total RNA was extracted from heart tissue using the Allprep DNA/RNA Mini Kit (Qiagen) per manufacturer protocol. Libraries were generated using the Illumina TruSeq Stranded mRNA Sample Prep Kit following manufacturer instructions (Illumina). 75 base pair single-end sequencing was preformed using the Illumina HiSeq 2500 platform. Read quality was assessed using FastQC, version 0.11.5 (Babraham Bioinformatics, Cambridge, UK). Reads were mapped to the mm10 genome using STAR v2.5.3a^34^. Gene expression levels were quantified across all exons using HOMER v4.10^35^. Differential gene expression analysis was carried out via edgeR v3.26.7^36,37^. Results corrected for multiple hypotheses testing using the Benjamini-Hochberg method^38^. The FDR threshold for significance was set at ≤0.05 and a log2 fold-change ≥1. Heatmaps and clustering were performed using R 3.6.1 (R Core Team) using variance stabilizing transformed counts from DESeq2 v1.24.0. Graphical packages (pheatmap v1.0.12, ggplot2 v3.3.2, gplots v3.0.1.1, RColorBrewer v1.1-2, VennDiagram v1.6.20) were used to visualize data by hierarchical clustering, and generating heat maps, Venn diagrams and PCA plots. GO and KEGG analyses were done using DAVID^39,40^.

### Statistics and data presentation

Prism 9 was used for graphing of data and statistical analyses. For survival, Log-rank analysis was used. D’Agostino and Shapiro-Wilk normality tests were performed on data sets to determine the distribution of the data sets. For pairwise comparisons, unpaired t-test, Mann-Whitney test, One-way ANOVA or Kruskal-Wallis test with Two-stage linear step-up procedure of Benjamini, Krieger and Yekutieli test, or Two-way ANOVA. Statistical tests used are noted in the figure legends. In accordance with our IACUC guidelines, in an effort to reduce the number of animals used for our experiments, where possible and appropriate, experiments were done in parallel and shared controls were used. This is indicated in the Figure legends where relevant. For some experiments, data are shown in multiple panels to simplify viewing of different comparisons within the data sets. This is indicated in the figure legends and source data where relevant.

## Acknowledgements

This work was supported by an NIH Pioneer award (DP1 AI144249) (JSA), a Keck Foundation Distinguished Young Scholar Award (JSA), NIH grant R01AI114929 (JSA), a Canadian Institutes of Health Research Postdoctoral Fellowship (JLM), the NOMIS Foundation, and the Razavi Newman Integrative Genomics and Bioinformatics Core Facility of the Salk Institute with funding from NIH-NCI CCSG: P30 014195, and the Helmsley Trust. Author contributions: JSA is responsible for conceptualization, performed experiments, analyzed data and wrote the paper; KSK performed in vivo experiments, troponin, wrote materials and methods; JLM performed in vivo experiments, analyzed data, wrote materials and methods; AEW performed all bioinformatics analyses and wrote materials and methods for RNAseq analysis; SJS performed qPCRs and western blots, wrote materials and methods. JMS developed histopathology scoring scheme and performed analyses and imaging. Special thanks to A. Ayres for help with the heart images.

