## Supplemental Figures and Legends for "Age-dependent roles of cardiac remodeling in sepsis defense and pathogenesis"

#### Supplemental Figure Legends

**Supplemental Figure 1. Original blot images**
(A-D) Original blots for Supplemental Figure 9A. (E-H) Original blots for Supplemental Figure 9B. (I-L) Original blots for Supplemental Figure 9G – young. (M-P) Original blots for Supplemental Figure 9G – old. The same lysates were run on two different gels in parallel. After transfer to membranes, the membranes were cut and probed for pFoxO1, GAPDH or total FoxO1.

**Supplemental Figure 2. Old and young mice exhibit differences in homeostatic variables.** Homeostatic variables of 12-week old and 75-week old uninfected mice were measured. (A) Blood pressure (B) Percent body fat (C) Percent lean body mass (D) Body weight (E) Percent water weight (F) Average daily food intake (G) Weights of organs (H) glucose tolerance test (I) Area under the curve analysis for (H) (J) Fasting glucose levels (K) Body temperature (L) Total bicarbonate in serum (M) Serum potassium levels (N) anion gap (O) VO2 and (P) area under the curve analysis for (O). n = 5 mice per age group. unpaired t-test. Error bars represent +/- SEM. * $p < 0.05$, ** $p < 0.005$, *** $p < 0.001$; **** $p < 0.0001$.

**Supplemental Figure 3**. **Body composition is not a predictor of outcome in old and young septic mice.** MRIs were taken of 12 and 75 week old mice. Following MRIs the mice were challenged with the $LD_{50}$ dose of polymicrobial sepsis and outcome was determined. (A) Starting body weight, (B) Body fat mass, (C) Percent body fat, (D) Lean body mass, (E) Percent lean body mass, (F) Total body water mass, (G) Percent total body water content. n = 4-6 mice per condition. Error bars indicate +/- SEM.

**Supplemental Figure 4. Thermoneutrality does not change susceptibility to polymicrobial sepsis in young and old mice**. 12 and 75-week old mice housed at 22°C or 30°C were challenged with polymicrobial sepsis. (A) Minimal body temperature of young mice exhibited during infection. n = 4-16 mice per group. ****$p$ 0.0001. (B) Minimal body temperature of old mice exhibited during infection. n = 4-6 mice per group. *$p < 0.05$, ****$p$ 0.0001. (C) Most severe morbidity score of young mice exhibited during infection. n = 4-16 mice per group. **$p < 0.01$. (D) Most severe morbidity score of old mice exhibited during infection. n = 4-6 mice per group. *$p < 0.05$, **$p < 0.01$. (E) Health trajectories of young mice exhibited during infection. n = 10-20 mice per condition. (F) Individual data points for (E). (G) Minimal health scores exhibited by young mice during infection. n = 4-16 mice per group. **$p < 0.005$, ***$p < 0.0005$. (H) Health trajectories of old mice exhibited during infection. n = 10-20 mice per condition. n = 10 mice per condition. (I) Individual data points for (H). (J) Minimal health scores exhibited by old mice during infection. n = 4-6 mice per group. **$p < 0.01$. (K) Survival of young mice. n = 10-20 mice per condition. $p = 0.3795$. (L) Survival of old mice. n = 10 mice per condition. $p = 0.8138$. Error bars indicate +/- SEM. Log rank analysis for survival. One-way ANOVA or Kruskal-Wallis with post Benjamini, Krieger and Yekutieli post-test for pairwise comparisons. ND indicates not determined because all mice were dead.

**Supplemental Figure 5. Young and old survivors and dying mice challenged with polymicrobial sepsis differ in disease tolerance**. 12 and 75-week old mice were challenged with the $LD_{50}$ dose of polymicrobial sepsis. Organs were harvested and levels of *E. coli* colonizing the organs were quantified. (A) Total *E. coli* CFUs in liver (B) Total *E. coli* CFUs in spleen (C) Total *E. coli* CFUs in kidney (D) Total *E. coli* CFUs in heart (I) Total *E. coli* CFUs in lung. n = 4-6 mice per condition. Geometric mean +/- SD and One-way ANOVA or Kruskal-Wallis with post Benjamini, Krieger and Yekutieli post-test for pairwise comparisons. ND indicates not determined because mice in that group were dead at that timepoint.

**Supplemental Figure 6. Young and old survivors and dying mice challenged with polymicrobial sepsis differ in disease tolerance**. 12 and 75-week old mice were challenged with the $LD_{50}$ dose of polymicrobial sepsis. Organs were harvested and levels of *S. aureus* colonizing the organs were quantified. (A) Total *S. aureus* CFUs in liver (B) Total *S. aureus* CFUs in spleen (C) Total *S. aureus* CFUs in kidney (D) Total *S. aureus* CFUs in heart (E) Total *S. aureus* CFUs in lung. n = 4-6 mice per condition. Geometric mean +/- SD and One-way ANOVA or Kruskal-Wallis with post Benjamini, Krieger and Yekutieli post-test for pairwise comparisons. ND indicates not determined because mice in that group were dead at that timepoint.

**Supplemental Figure 7. Age dependent health and disease trajectories exhibited by polymicrobial sepsis $LD_{50}$ challenged mice**. (A-C) 12-week and 75-week old mice were challenged with the $LD_{50}$ dose of polymicrobial sepsis. Temperature, and morbidity were measured to calculate health scores. (A) The minimal body temperature exhibited by mice. n=4-5 mice per condition, ****$p<0.0001$, **$p<0.005$, *$p<0.05$. (B) The most severe morbidity score exhibited by mice (lower score indicates greater morbidity). n = 4-5 mice per condition, **$p<0.005$. (C) Minimal health scores exhibited by mice over the course of the infection. n = 4-5 mice per condition, **$p<0.005$, *$p<0.08$. (D-F)12-week old mice were infected with the $LD_{50}$ dose of polymicrobial sepsis and (D) Temperature and (E) Morbidity were determined every two hours to calculate the (F) Health scores over time. n = 5 mice per condition, *$p < 0.05$, **$p < 0.005$; ***$p < 0.0005$; ****$p < 0.0001$. Data in (F) are the same as those in Figure 2B and are showing the individual data points. Minimal temperature, morbidity and health scores shown in (A-C) were obtained from the time courses shown in (D-F). (G-I) 75-week old mice were infected with the $LD_{50}$ dose of polymicrobial sepsis and (G) Temperature and (H) Morbidity were determined every two hours post-infection to calculate the (I) Health scores over time. n = 5 mice per condition, * $p < 0.05$, ** $p < 0.005$; *** $p < 0.0005$; $p < 0.0001$. Data in (I) are the same as those in Figure 2C and are showing the individual data points. Minimal temperature, morbidity and health scores shown in (A-C) were obtained from the time courses shown in (D-I). Error bars indicate +/-SEM. For pairwise comparisons, unpaired t-test, Mann-Whitney test, One-way ANOVA or Kruskal-Wallis test with Two-stage linear step-up procedure of Benjamin, Krieger and Yekutieli, or Two-way ANOVA.

**Supplemental Figure 8. Age dependent health and disease phenotypes exhibited by polymicrobial sepsis $LD_{50}$ challenged mice**. (A-C) 12-week and 75-week old mice were challenged with the $LD_{50}$ dose of polymicrobial sepsis and markers of organ damage were measured (A) BUN, (B) AST and (C) ALT. Young uninfected n = 18-23, young surviving n = 20-21, young dying n = 25-27, old uninfected n = 7-9, old surviving n = 10, old dying n = 16-19. *$p < 0.05$, **$p < 0.01$; ***$p < 0.005$, ****$p < 0.0001$. (D) Original, unedited images from Figure 2F. (E-G) Young and old mice were challenged with the $LD_{50}$ dose of polymicrobial sepsis. When dying mice for each age group reached minimal health (clinical endpoint), hearts were harvested from age-matched infected surviving and uninfected control mice and RNAseq was done (10 hrs post-infection for young mice and 24 hrs for old mice). (E) PCA plot of normalized raw cardiac counts. (F) Venn diagrams of genes that were depleted in young dying hearts compared to both young surviving and young uninfected mice vs old dying hearts compared to both old surviving and old uninfected mice, and young surviving hearts compared to both young dying and young uninfected mice vs old surviving hearts compared to both old dying and old uninfected mice. (G) Gene ontology analysis was done on the distinct gene lists for each group as shown in the bar graphs. (H) Serum Troponin I levels of $LD_{50}$-challenged and uninfected age matched mice. n = 8-11 mice per condition. *** $p < 0.0005$. (I) Serum Galectin-3 levels of $LD_{50}$-challenged and uninfected age matched mice. n = 9-18 mice per condition. * $p < 0.05$, ** $p < 0.005$; *** $p < 0.0005$. (J) Young and old mice were challenged with the $LD_{50}$ dose of polymicrobial sepsis and organs were harvested for histopathology analysis. Representative images of spleen (left column, bar = 100 microns); lung (2$^{nd}$ column from left, bar = 50 microns); liver (2$^{nd}$ column from right, bar = 50 microns); and kidney (right column, bar = 100 microns) in (from top to bottom) control uninfected young mice; infected surviving young mice; infected dying young mice; control uninfected old mice; infected surviving old mice; and infected dying old mice.  Compared to other groups, infected dying young and infected dying old mice have expanded splenic red pulp (congestion, asterisks); increased congestion and leukocyte infiltration in the lung interstitium (arrows); increased sinusoidal congestion in the liver (arrows); and increased congestion in the kidney medulla (asterisks). In the lung, some old mice have perivascular and peribronchiolar lymphoid aggregates (asterisk) as an incidental finding. Error bars indicate +/-SEM. For pairwise comparisons, unpaired t test, Mann-Whitney test, One-way ANOVA or Kruskal-Wallis test with Two-stage linear step-up procedure of Benjamin, Krieger and Yekutieli, or Two-way ANOVA.

**Supplemental Figure 9. Age dependent regulation of cardiac FoxO1 during sepsis.** (A-B) Levels of total FoxO1 and active FoxO1 in hearts from old and young uninfected mice under fed and fasted conditions. (A) Western blot of total FoxO1, pFoxO1 and GAPDH under fasted conditions. (B) Western blot of total FoxO1, pFoxO1 and GAPDH under fed conditions. n = 5 mice per condition. (C) Cardiac expression of *Foxo1* in hearts from old and young mice under fed and fasted conditions. n = 5 mice per condition. (D) Expression of *Foxo1* in uninfected and $LD_{50}$-challneged old and young mice at 10hrs (young and old) and 24hrs post-infection. n = 3-5 mice per condition. ****$p < 0.0001$. Heart data are also shown in Figure 3D. (E) Western blots of total FoxO1 and active FoxO1 in hearts from old and young uninfected and $LD_{50}$-challenged mice when dying animals reached maximal morbidity for each age group (10hrs for young and 24hrs for old). n = 3 mice per condition.

Error bars indicate +/- SEM. For pairwise comparisons, One-way ANOVA or Kurskal Wallis with Two-stage linear step-up procedure of Benjamini, Krieger and Yekutieli.

**Supplemental Figure 10. FoxO1 protects young hosts from sepsis induced morbidity and mortality.** (A-C) Uninfected and polymicrobial sepsis infected young mice were treated with a vehicle or FoxO1 inhibitor. (A) Minimal body temperature, (B) Most severe morbidity score and (C) health trajectories exhibited by vehicle surviving, FoxO1 inhibitor dying during infection and uninfected mice. n = 5-10 mice per condition. *$p < 0.05$, **$p < 0.005$, ***$p < 0.0005$, ****$p < 0.0001$. (D-H) Young and old *Foxo1 mck$^{cre-}$* and *mck$^{cre+}$* mice were challenged with polymicrobial sepsis. (D) Minimal body temperature during infection, (E) Body temperature just prior to infection, (F) Most severe morbidity score, (G) Morbidity scores just before infection and (H) Health trajectories during infection. n = 6-11 mice per condition. *$p < 0.05$, **$p < 0.01$, ***$p < 0.005$, ****$p < 0.0001$. (I-K) Colony forming units (CFU) in shown organs/tissues of uninfected and infected young vehicle-treated and FoxO1 inhibitor-treated mice. (I) Total CFUs, (J) *E. coli* CFUs, (K) *S. aureus* CFUs. n = 4-5 mice per condition. For samples that had CFU counts below the limit of detection they are noted on the graph as “X BLD” where “X” indicates the number of mice that had counts BLD for that organ/tissue. (L-M) Serum cytokines of uninfected and infected young vehicle-treated and FoxO1 inhibitor-treated mice. (L) Heat map of infected conditions normalized to uninfected conditions. (M) Individual data points. n = 4-5 mice per condition. Vehicle U – vehicle uninfected, FoxO1 U – FoxO1 inhibitor uninfected, Vehicle I – vehicle infected, FoxO1 I – FoxO1 inhibitor infected. Error bars indicate +/- SEM. For CFU plots, geometric mean +/- SD. For pairwise comparisons, unpaired t-test, Mann-Whitney test, One-way ANOVA or Kurskal Wallis with Two-stage linear step-up procedure of Benjamini, Krieger and Yekutieli, or Two-way ANOVA.

**Supplemental Figure 11. FoxO1 protects young hosts from sepsis induced cardiomegaly and heart failure.** (A) Original images of hearts shown in Figure 3G. (B-K) Histopathology analysis of vehicle treated surviving and FoxO1 inhibitor treated dying infected mice, and uninfected controls at 10hrs post-infection. (B) Heart edema score, (C) Cardiomyocyte score, (D) Heart leukocyte score, (E) Heart congestion score, (F) From top to bottom, representative images of heart (left ventricle) in vehicle uninfected, inhibitor uninfected, vehicle infected, and inhibitor infected animals. Compared to the other groups, the inhibitor infected animals have increased numbers of vessels expanded by red blood cells (congestion) and leukocytes (arrows) with mild edema separating the cardiomyocytes. The ventricular myocardium wall of the inhibitor infected animal also has regions of cardiomyocyte pallor and vacuolation (asterisk) and slightly enlarged, hypereosinophilic cardiomyocytes (large arrow). Bar = 100 microns. (G) Liver congestion score, (H) Spleen congestion score, (I) Lung congestion score, (J) Kidney congestion score, (K) Representative images of spleen (top row, bar = 100 microns); lung (2$^{nd}$ row from top, bar = 50 microns); liver (2$^{nd}$ from bottom, bar = 50 microns); and kidney (bottom row, bar = 100 microns) in vehicle uninfected (left column), inhibitor uninfected (2$^{nd}$ from left), vehicle infected (2$^{nd}$ from right), and inhibitor infected (right column) mice. Compared to other groups, inhibitor infected mice have expanded splenic red pulp (congestion, asterisk); increased congestion and leukocyte infiltration in the lung interstitium (small arrows) and increased leukocytes within pulmonary vessels (large arrow); increased sinusoidal congestion in the liver (arrows); and increased congestion in the kidney medulla (arrows). n = 5-17 mice per condition. *$p < 0.05$, **$p < 0.005$, ***$p < 0.0005$, ****$p < 0.0001$. (L-N) Serum levels of organ damage markers from uninfected and infected young vehicle and FoxO1 inhibitor treated mice. (L) AST, (M) ALT, (N) BUN. 8-23. **$p < 0.005$, ***$p < 0.005$, ****$p < 0.0001$. Vehicle U – vehicle uninfected, FoxO1 U – FoxO1 inhibitor uninfected, Vehicle I – vehicle infected, FoxO1 I – FoxO1 inhibitor infected. Error bars indicate +/- SEM. For pairwise comparisons, One-way ANOVA or Kurskal Wallis with Two-stage linear step-up procedure of Benjamini, Krieger and Yekutieli, or Two-way ANOVA.

**Supplemental Figure 12**. **FoxO1 regulation of Trim63 is cardioprotective in young septic mice**. (A-B) Organs and tissues from uninfected 12 and 75-week old mice under fed and fasted conditions were harvested and the expression of (A) *Trim63* and (B) *Fbxo32* were measured. n = 5 mice per condition. *$p < 0.05$. (C-D) 12 and 75-week old mice were infected with the $LD_{50}$ dose of polymicrobial sepsis. Hearts were harvested at 10hrs (young and old) and at 24hrs (old). Expression of (C) *Trim63* and (D) *Fbxo32* were measured. n = 3-5 mice per condition. *$p < 0.05$, ***$p < 0.0005$, ****$p < 0.0001$. Data are also presented in **Figure 4B-C**. (E-G) Young wild type and *Trim63* $^{+/-}$ mice were infected with a low dose of polymicrobial sepsis (~$LD_{25}$) and (E) Minimal temperature exhibited in the first 10hrs post-infection, (F) Most severe morbidity score exhibited in the first 10hrs post-infection and health trajectories were determined. n = 5-9 mice per condition. *$p < 0.05$, **$p < 0.01$, ***$p < 0.0005$. (H) Original images of hearts shown in Figure 4G. (I-L) Young wild type and *Trim63* $^{+/-, -/-}$

mice were infected with a low dose of polymicrobial sepsis (~$LD_{25}$) and organs were harvested for histopathology analysis. (I) Histologic images of heart (left ventricle). Compared to the other groups, the *Trim63*$^{+/-,-/-}$ infected animals have increased myocardial congestion (arrows) and, in this case, focal congestion with hemorrhage (asterisk). The ventricular myocardium wall of the KO infected animal also has regions of cardiomyocyte pallor and vacuolation (asterisk) and slightly enlarged, hypereosinophilic cardiomyocytes with vessels containing mildly increased numbers of leukocytes (arrow). Bar = 100 microns. (J) Heart edema score; (K) Heart leukocyte score; (L) Cardiomyocyte score. n = 10-16 mice per condition. $^*p < 0.05$; $^{**}p < 0.005$, $^{***}p < 0.0005$, $^{****}p < 0.0001$. *Trim*$^{+/+}$ U = wild type uninfected; *Trim63*$^{+/-,-/-}$ U = heterozygous and homozygous Trim63 mutants uninfected; *Trim*$^{+/+}$ I = wild type infected; *Trim63*$^{+/-,-/-}$ I = heterozygous and homozygous Trim63 mutants infected. Error bars indicate +/- SEM. For pairwise comparisons, Mann-Whitney test, One way ANOVA, Kurskal Wallis with Two-stage linear step-up procedure of Benjamini, Krieger and Yekutieli, Two-way ANOVA.

**Supplemental Figure 13**. **FoxO1 and Trim63 do not regulate skeletal muscle mass in young septic mice.** (A-F) Young *Trim63*$^{+/+}$ and *Trim63*$^{+/-,-/-}$ mice were infected with polymicrobial sepsis or left uninfected. Organs were harvested at 10hrs post-infection for histopathology analysis. (A) Representative images of spleen (top row, bar = 100 microns); lung (2$^{nd}$ row from top, bar = 50 microns); liver (2$^{nd}$ from bottom, bar = 50 microns); and kidney (bottom row, bar = 100 microns) in WT uninfected (left column), *Trim63*$^{+/-,-/-}$ uninfected (2$^{nd}$ from left), WT infected (2$^{nd}$ from right), and *Trim63*$^{+/-,-/-}$ infected (right column) mice. Compared to other groups, *Trim63*$^{+/-,-/-}$ infected mice have expanded splenic red pulp (congestion, asterisk); increased congestion and leukocyte infiltration in the lung interstitium (small arrows) and increased leukocytes within pulmonary vessels (large arrow); increased sinusoidal congestion in the liver (arrows); and increased congestion in the kidney medulla (arrows). Semi-quantitative scoring of (B) Cardiac congestion, (C) Liver congestion, (D) Spleen congestion, (E) Kidney congestion, (F) Lung congestion. n = 10-16 mice per condition. $^*p < 0.05$, $^{**}p < 0.01$, $^{***}p < 0.0005$, $^{****}p < 0.0001$. (G-H) 12 and 75 week old mice were infected with polymicrobial sepsis. The (G) Tibialis anterior (TA) and (H) Extensor digitorum longus (EDL) leg muscles were harvested when dying animals in each age group each maximal morbidity (10 hrs for young and 24 hrs for old) and expression of *Trim63* was measured. n = 6-10 mice per condition. $^{**}p < 0.005$, $^{****}p < 0.0001$. (I-J) Young mice were infected with a low dose (~$LD_{10}$) of polymicrobial sepsis and treated with a FoxO1 inhibitor or vehicle. The (I) TA and (J) EDL were harvested at 10 hr post-infection and expression of *Trim63* was measured and normalized to *Rps17*. n = 4-9 mice per condition. $^{**}p < 0.01$. (K-N) Young *Trim63*$^{+/+}$ and *Trim63*$^{+/-,-/-}$ mice were infected with polymicrobial sepsis or left uninfected. The (K) Quad; (L) TA; (M) EDL and (N) Gas were dissected at 10hrs post-infection and normalized to body weight. n = 8-15 mice per condition. (O-R) Leg muscles were dissected from uninfected and $LD_{50}$-challenged old and young mice when dying mice reached maximal morbidity (10hrs for young and 24hrs for old). (O) Quad; (P) TA; (Q) EDL and (R) Gas. n = 6-10 mice per condition. $^*p < 0.05$, $^{**}p < 0.01$, $^{***}p < 0.0005$. (S-U) Serum levels of (S) AST, (T) ALT and (U) BUN of young uninfected and infected *Trim63*$^{+/+}$ and *Trim63*$^{+/-,-/-}$ mice at 10hrs post-infection. n = 7-11 mice per condition. $^*p < 0.05$, $^{**}p < 0.01$, $^{***}p < 0.0005$, $^{****}p < 0.0001$. Vehicle U – uninfected vehicle treated, FoxO1 U – uninfected FoxO1 inhibitor treated, Vehicle I – vehicle treated infected, FoxO1 I – FoxO1 inhibitor treated infected. *Trim*$^{+/+}$ U = wild type uninfected; *Trim63*$^{+/-,-/-}$ U = heterozygous and homozygous Trim63 mutants uninfected; *Trim*$^{+/+}$ I = wild type infected; *Trim63*$^{+/-,-/-}$ I = heterozygous and homozygous Trim63 mutants infected. Error bars indicate +/- SEM. For pairwise comparisons, Mann-Whitney test, One way ANOVA, Kurskal Wallis with Two-stage linear step-up procedure of Benjamini, Krieger and Yekutieli, Two-way ANOVA.

**Supplemental Figure 14**. **FoxO1 regulation of Trim63 is a driver of sepsis pathogenesis and cardiac cachexia in old septic mice**. (A) 75-week old mice were infected with polymicrobial sepsis and treated with a FoxO1 inhibitor or vehicle. Survival was monitored. n = 5 mice per condition. (B-D) 75-week old mice were treated with a FoxO1 inhibitor or vehicle and infected with polymicrobial sepsis. (B) Minimal body temperature exhibited during infection, (C) Most severe morbidity score observed during infection, (D) Health trajectories exhibited during infection. Values for uninfected are the values exhibited by mice just prior to infection. n = 10 mice per condition. $^{**}p$ 0.005, $^{****}p < 0.0001$. (E) Representative heart images from uninfected and infected old mice treated with a FoxO1 inhibitor or vehicle control. (F) Original images of hearts shown in (E) and Figure 4N. (G) Heart weights normalized to body weights of uninfected and infected old mice treated with vehicle, FoxO1 or Trim63 inhibitor at 24hrs post-infection. Vehicle and Trim63 values also shown in Figure 4O.(H) Values from (G) normalized to the average of Vehicle U values from (G). n = 5-23 mice per condition. $^*p < 0.05$, $^{**}p < 0.005$, $^{***}p < 0.0005$, $^{****}p < 0.0001$. Vehicle and Trim63 values also shown in Figure 4P. (I)

Cardiac expression of *Trim63* in uninfected and infected old mice treated with FoxO1 inhibitor or vehicle control at 10hrs post-infection. n = 4-10 mice per condition. *$p < 0.05$, **$p < 0.01$, ****$p < 0.0001$. (J-L) 72-week old $Trim63^{+/+}$ and $Trim63^{-/-}$ mice were infected with polymicrobial sepsis. Survival, temperature and morbidity were monitored. (J) Survival; (K) Minimal temperature exhibited by dying $Trim63^{+/+}$ and surviving $Trim63^{-/-}$ mice within first 24hrs of infection; (L) Most severe morbidity score exhibited by dying $Trim63^{+/+}$ and surviving $Trim63^{-/-}$ mice within first 24hrs of infection; (M) Health trajectories of dying $Trim63^{+/+}$ and surviving $Trim63^{-/-}$ mice within first 24hrs of infection. n = 2-5 mice per condition. *$p < 0.05$, **$p < 0.01$, ***$p < 0.0005$. Uninfected values were values obtained from mice at t=0hr just prior to infection. (N-P) 75-week old mice were infected with polymicrobial sepsis and treated with a Trim63 inhibitor or vehicle. (N) minimal temperature exhibited by vehicle treated dying and Trim63 inhibitor treated surviving mice within first 24hrs of infection; (O) Most severe morbidity score exhibited by vehicle treated dying and Trim63 inhibitor treated surviving mice within first 24hrs of infection; (P) Health trajectories of vehicle treated dying and Trim63 inhibitor treated surviving mice within first 24hrs of infection. n = 6 mice per condition. *$p < 0.05$, **$p < 0.005$, ***$p < 0.0005$, ****$p < 0.0001$. Uninfected values were values obtained from mice at t=0hr just prior to infection. (Q-T) 75-week old mice were infected with polymicrobial sepsis or left uninfected and treated with a Trim63 inhibitor or vehicle. At 24 hrs post-infection hearts were harvested. (Q) Heart edema score, (R) Cardiomyocyte score, (S) Heart leukocyte infiltration score. (T) From left to right, representation images of heart (left ventricle) in vehicle uninfected, Trim63 inhibitor uninfected, vehicle infected, and Trim63 inhibitor infected animals. Compared to the other groups, the vehicle infected animals have increased numbers of vessels expanded by red blood cells (congestion, asterisk) and leukocytes (thin arrow) and mild edema. Bacteria are observed in this animal (large arrow). The ventricular myocardium of the vehicle infected animal also has regions of cardiomyocyte pallor and vacuolation and slightly enlarged, hypereosinophilic cardiomyocytes (within circle). Bar = 100 microns. n = 10-15 mice per condition. **$p < 0.005$, ***$p < 0.0005$, ****$p < 0.0001$. Vehicle U – uninfected vehicle treated, FoxO1 U – uninfected FoxO1 inhibitor treated, Vehicle I – vehicle treated infected, FoxO1 I – FoxO1 inhibitor treated infected. Trim63 U – Trim63 inhibitor uninfected, Trim63 I – Trim63 inhibitor infected. $Trim^{+/+}$ U = wild type uninfected; $Trim63^{+/-, -/-}$ U = heterozygous and homozygous Trim63 mutants uninfected; $Trim^{+/+}$ I = wild type infected; $Trim63^{+/-, -/-}$ I = heterozygous and homozygous Trim63 mutants infected. Error bars indicate +/- SEM. For pairwise comparisons, One way ANOVA, Kurskal Wallis with Two-stage linear step-up procedure of Benjamini, Krieger and Yekutieli, Two-way ANOVA. For survival, Log-rank analysis.

**Supplemental Figure 15**. **FoxO1 regulation of Trim63 is a driver of sepsis induced cardiac failure in old septic mice**. (A) 75-week old mice were infected with polymicrobial sepsis and treated with a Trim63 inhibitor or vehicle. Serum Galectin-3 was measured. n = 10-14 mice per condition. ***p < 0.0005, ****p < 0.0001. (B) 75-week old mice were infected with polymicrobial sepsis and treated with a Trim63 inhibitor or vehicle. Serum Troponin I was measured. n = 4-5 mice per condition. (C-H) 75-week old mice were infected with polymicrobial sepsis or left uninfected and treated with a Trim63 inhibitor or vehicle. At 24 hrs post-infection organs were harvested for histopathology analysis. (C) Representative images of spleen (top row, bar = 100 microns), liver (middle row, bar = 50 microns), and kidney (bottom row, bar = 50 microns) in vehicle uninfected (left column), Trim63 inhibitor uninfected (2nd from left), vehicle infected (2nd from right), and Trim63 inhibitor infected (right column) mice. Compared to other groups, vehicle infected mice have expanded splenic red pulp (congestion, asterisk); increased sinusoidal congestion and areas of hemorrhage (asterisk) in the liver; and acute necrosis of the tubules in the kidney (arrows). Semi-quantitative scoring of (D) heart congestion, (E) liver congestion, (F) Spleen congestion, (G) kidney congestion, (H) lung congestion. n = 10-15 mice per condition. *$p < 0.05$, **$p < 0.005$, ****$p < 0.0001$. (I-K) Serum levels of (I) AST, (J) ALT and (K) BUN of old uninfected and infected mice treated with Trim63 inhibitor or vehicle. n = 5-12 mice per condition. *$p < 0.05$, **$p < 0.01$, ***$p < 0.005$. (L-O) 75 week old mice were infected with polymicrobial sepsis and treated with a Trim63 inhibitor or vehicle. Leg muscles were dissected, weighed and normalized to body weight. (L) quad; (M) TA; (N) EDL and (O) Gas. . n = 10-13 mice per condition. *$p < 0.05$; **$p < 0.01$. (P-Q) 75-week old mice were infected with polymicrobial sepsis and treated with a FoxO1 inhibitor or vehicle. *Trim63* expression was measured in the (P) TA and (Q) EDL. n = 5-10 mice per condition. *$p < 0.05$; **$p < 0.01$, ***$p < 0.005$, ****$p < 0.0001$. Vehicle U – uninfected vehicle treated, FoxO1 U – uninfected FoxO1 inhibitor treated, Vehicle I – vehicle treated infected, FoxO1 I – FoxO1 inhibitor treated infected. Trim63 U – Trim63 inhibitor uninfected, Trim63 I – Trim63 inhibitor infected. Error bars indicate +/- SEM. For pairwise comparisons, One way ANOVA, Kurskal Wallis with Two-stage linear step-up procedure of Benjamini, Krieger and Yekutieli.

**KEY RESOURCES TABLE**

| REAGENT or RESOURCE | SOURCE | IDENTIFIER |
|---|---|---|
| Antibodies, inhibitors | | |
| FoxO1 inhibitor AS1842856 (powder) 10mg | EMB Millipore | Cat# 344355-10MG |
| FoxO1 inhibitor in-solution AS1842856 5mg | EMB Millipore | Cat# 5.060810001 |
| TRIM63 EMBL inhibitor | Glixx Laboratories | Cat# GLXC-15396 |
| GAPDH (14C10) Rabbit mAb | Cell Signaling | 2118S |
| Phospho-FoxO1 (Thr24)/FoxO3a (Thr32) Antibody | Cell Signaling | 9464S |
| FoxO1 (C29H4) Rabbit mAb | Cell Signaling | 2880S |
| Anti-rabbit IgG HRP-linked Ab | Cell Signaling | 7074S |
| Protease Inhibitor Cocktail | Sigma | P2714 |
| Bacterial and virus strains | | |
| *E. coli* O21:H+ | Ayres et al., 2012 | |
| *S. aureus* (ATCC strain 12600) | ATCC | NCTC 8532 |
| Chemicals, peptides, and recombinant proteins | | |
| Neutral Buffered Formalin, 10% | Millipore | Cat# 65346-85 |
| D-(+)-Glucose | Sigma | Cat# G7021 |
| Ethanol 190 proof | Decon Laboratories, Inc | Cat# 2801 |

| | | |
|---|---|---|
| LB Media Components:<br><br>Bacto Tryptone<br><br>Bacto Yeast Extract<br><br>Sodium Chloride<br><br>Difco Agar Technical solidifying agent | <br><br>Gibco<br><br>Gibco<br><br>Fisher Chemical<br><br>BD | <br><br>Cat# 212750<br><br>Cat# 211705<br><br>Cat# S271-3<br><br>Cat# 281210 |
| Eosin Methylene Blue Agar (Levine) | Oxoid | Cat# CM0069 |
| Antibiotics for EMB Agar (AVNM):<br><br>Ampicillin Sodium Salt (1mg/ml)<br><br>Vancomycin hydrochloride (.5mg/ml)<br><br>Neomycin sulfate (1mg/ml)<br><br>Metronidazole (1mg/ml) | <br><br>Fisher Bioreagents<br><br>ACROS Organics<br><br>Fisher Bioreagents<br><br>MP Biomedicals | <br><br>Cat# BP1760-25<br><br>Cat# 1404-93-9<br><br>Cat# BP2669-25<br><br>Cat# 155710 |
| UltraPure distilled water | Invitrogen | Cat# 10977-015 |
| DPBS (1X)<br><br>[-] Calcium Chloride<br><br>[-] Magnesium Chloride | Gibco | Cat# 14190-144 |
| TBST:<br><br>Tris, Hydrochloride, ULTROL® Grade<br><br>Sodium Chloride<br><br>TWEEN® 20 | <br><br>Millipore Sigma<br><br>Fisher Chemical<br><br>Sigma-Aldrich | <br><br>1185-53-1<br><br>S2713<br><br>P1379-500ML |
| Trans-Blot Turbo Midi 0.2 µm Nitrocellulose Transfer Packs | Bio-Rad | 1704159 |
| Trans-Blot® Turbo™ Transfer System | Bio-Rad | 1704150 |
| Ponceau Total Protein Stain | Prometheus Protein Biology Products | 20-311 |
| Bovine Serum Albumin, Heat Shock Treated | Fisher BioReagents™ | BP1600100 |
| Prometheus OneBlock™ Western-CL Blocking Buffer | Prometheus Protein Biology Products | 20-313 |

| | | |
|---|---|---|
| ProSignal® Femto ECL Reagent | Prometheus Protein Biology Products | 20-302 |
| ProSignal® Dura ECL Reagent | Prometheus Protein Biology Products | 20-301 |
| Gel Doc XR+ Gel Documentation System | Bio-Rad | 1708195 |
| Allprep DNA/RNA Mini Kit | Qiagen | 80204 |
| SuperScript™ IV Reverse Transcriptase | Invitrogen | 18090010 |
| QuantStudio 5 Real-Time PCR instrument | Applied Biosystems | A28139 |
| Pierce™ BCA Protein Assay Kit | Thermo Scientific™ | 23225 |
| 2-Mercaptoethanol (50 mM) | Gibco™ | 31350010 |
| NuPAGE™ LDS Sample Buffer (4X) | Invitrogen™ | NP0007 |
| NuPAGE™ 7%, Tris-Acetate, 1.0 mm, Mini Protein Gel, 12-well | Invitrogen | EA03552BOX |
| NuPAGE™ Tris-Acetate SDS Running Buffer (20X) | Thermo Fisher | LA0041 |
| Dimethyl sulfoxide | Sigma | Cat# D8418-100ml |
| Tissue Extraction Reagent II | Invitrogen | FNN0081 |
| Critical commercial assays | | |
| Ultra-Sensitive Mouse Cardiac Troponin-I ELISA Kit | Life Diagnostics | Cat# CTNI-1-US |
| GAL3 ELISA Kit | Invitrogen | EMCGALS3 |
| BNP ELISA Kit | Cusabio | CSB-E07971m |
| Experimental models: Organisms/strains | | |
| C57BL/6 | Jackson Laboratories | 000664 |
| *Trim63*$^{-/-}$ | This study | |
| *Foxo1 mck cre* mice | Mark Febrraio, Monash University | |
| Oligonucleotides | | |

| Gene target | Forward 5’-3’ | Reverse 5’-3’ |
|---|---|---|
| *Rps17* | CGCCATTATCCCCAGCAAGA | CACAGGCCCTCTCTGAATCC |
| *FoxO1* | GGGTCCCACAGCAACGATG | CACCAGGGAATGCACGTCC |
| *Atrogin1* | CAGCTTCGTGAGCGACCTC | GGCAGTCGAGAAGTCCAGTC |
| *Murf* | AGTGTCCATGTCTGGAGGTCGTTT | ACTGGAGCACTCCTGCTTGTAGAT |
| Software and algorithms | | |
| Andrews, S. (2010). FastQC: A Quality Control Tool for High Throughput Sequence Data [Online]. Available online at: http://www.bioinformatics.babraham.ac.uk/projects/fastqc/ | | |
| R Core Team (2020). R: A language and environment for statistical computing. R Foundation for Statistical Computing, Vienna, Austria. URL https://www.R-project.org/. | | |
| Raivo Kolde (2019). pheatmap: Pretty Heatmaps. R package version 1.0.12. https://CRAN.R-project.org/package=pheatmap | | |
| H. Wickham. ggplot2: Elegant Graphics for Data Analysis. Springer-Verlag New York, 2016. | | |

| | | |
|---|---|---|
| Gregory R. Warnes, Ben Bolker, Lodewijk Bonebakker, Robert Gentleman, Wolfgang Huber Andy Liaw, Thomas Lumley, Martin Maechler, Arni Magnusson, Steffen Moeller, Marc Schwartz and Bill Venables (2019). gplots: Various R Programming Tools for Plotting Data. R package version 3.0.1.1. https://CRAN.R-project.org/package=gplots | | |
| Erich Neuwirth (2014). RColorBrewer: ColorBrewer Palettes. R package version 1.1-2. https://CRAN.R-project.org/package=RColorBrewer | | |
| Hanbo Chen (2018). VennDiagram: Generate High-Resolution Venn and Euler Plots. R package version 1.6.20. https://CRAN.R-project.org/package=VennDiagram | | |
| Other | | |
| NovaMax Plus blood glucose monitoring system | ADW Diabetes | Cat# 8548043435 |
| CLAMS | Columbus Instruments | |
| BD Microtainers SST | BD | Cat# 2022-04030 |
| Microvette® CB 300Z | | |
| CODA® Monitor Noninvasive Blood Pressure System | Kent Scientific Corporation | |
| Far Infrared Warming Platform | Kent Scientific Corporation | Product # AHP-405L |
| Digi-Sense Type J/K/T Thermocouple meter | Digi-Sense | Cat# 20250-91 |

Supplemental Figure 1_1

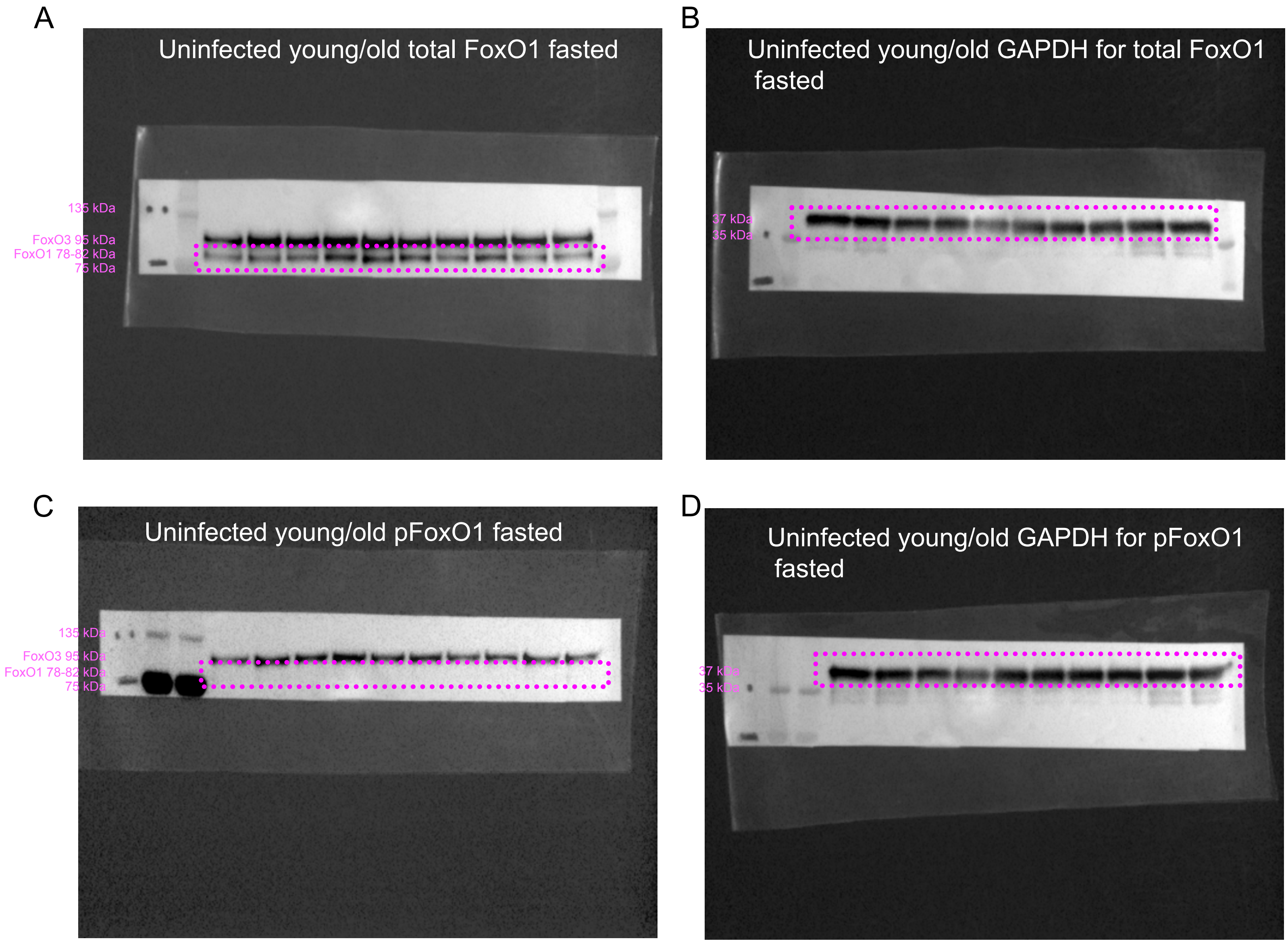

Supplemental Figure 1_2

E

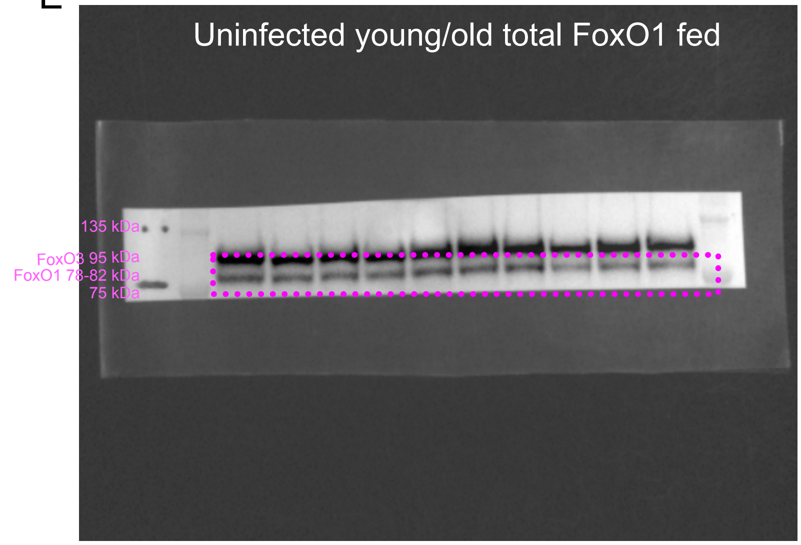

F

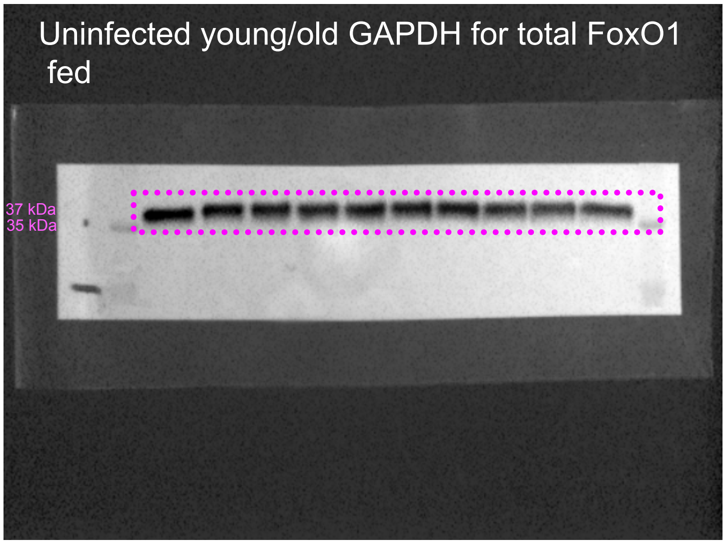

G

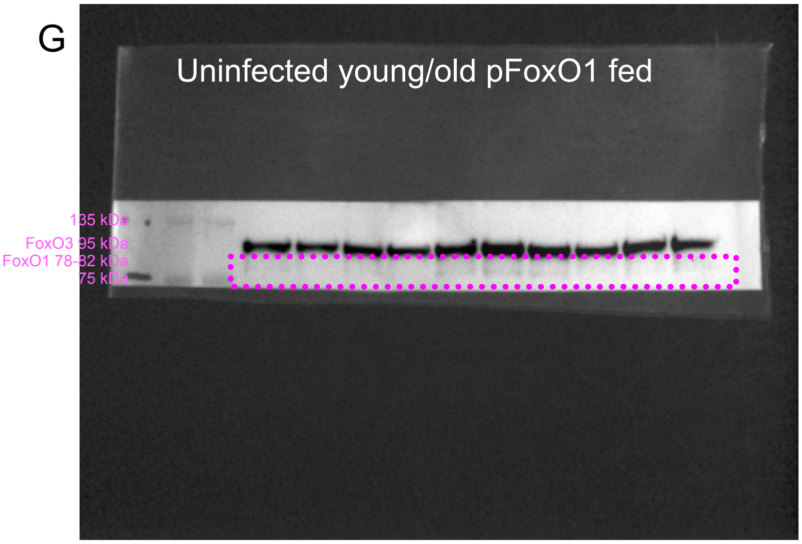

H

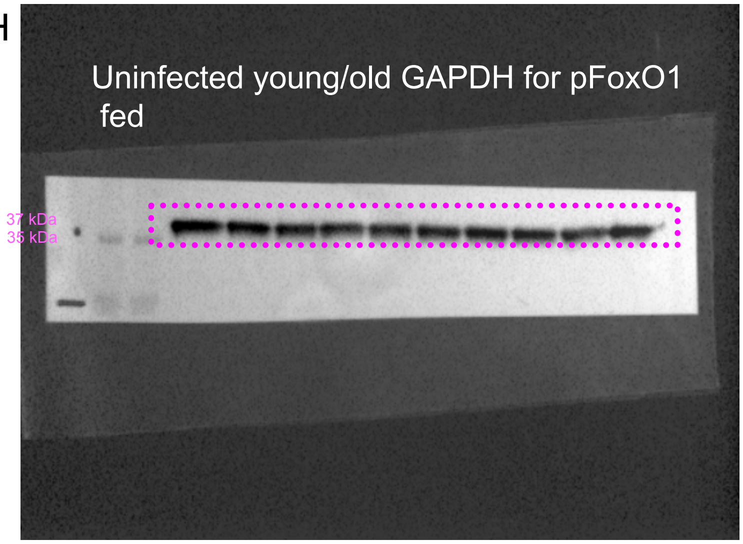

Supplemental Figure 1_3

I

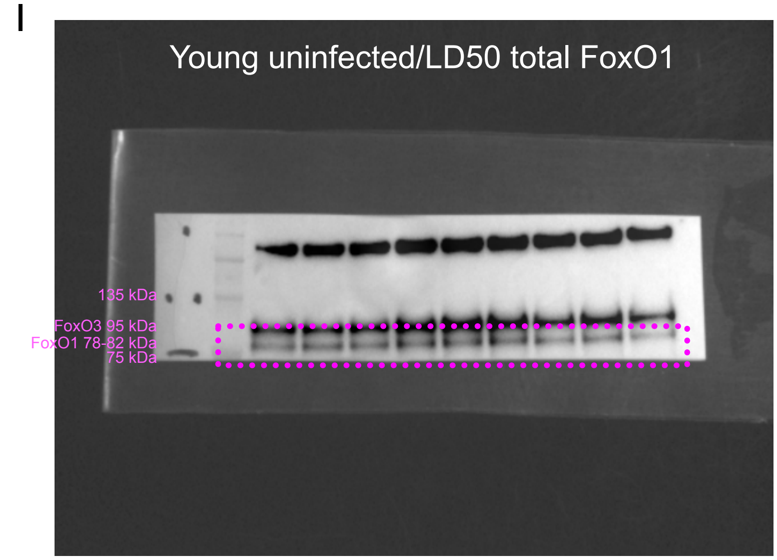

J

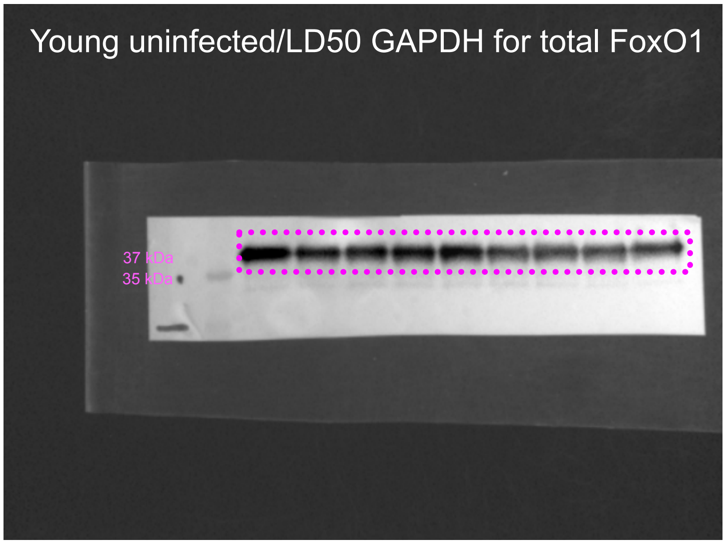

K

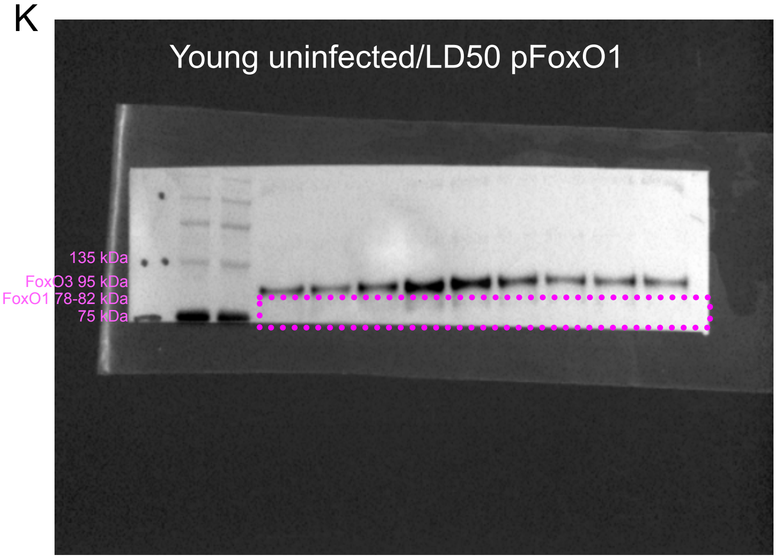

L

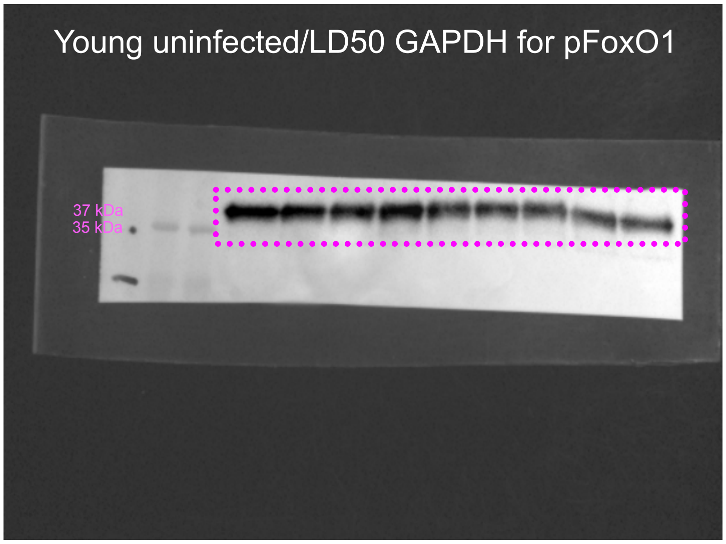

Supplemental Figure 1_4

M

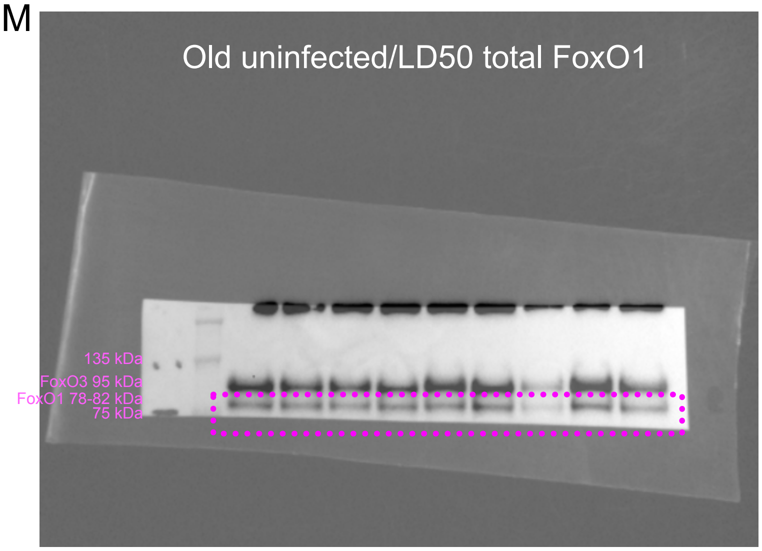

N

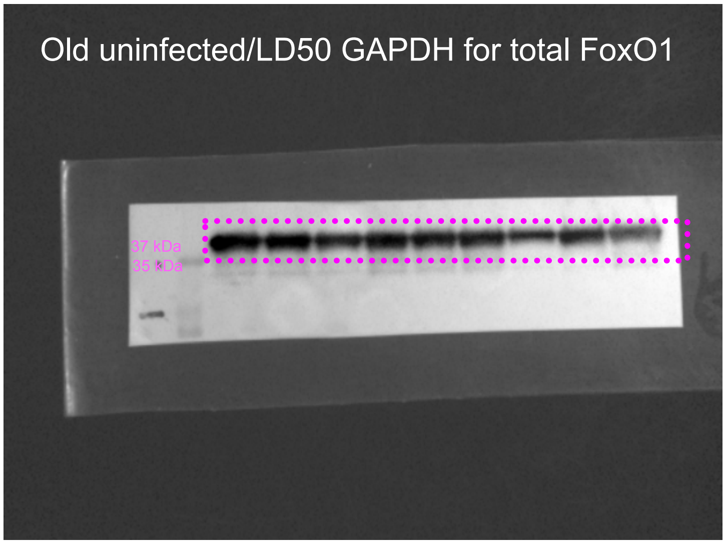

O

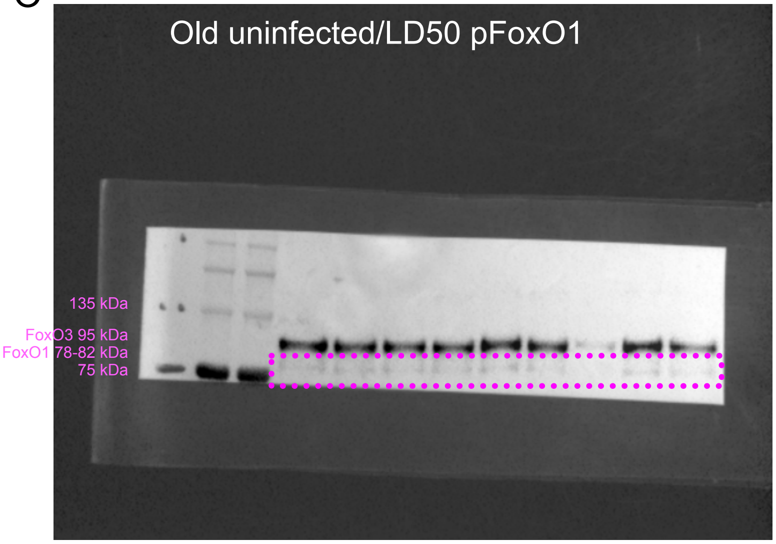

P

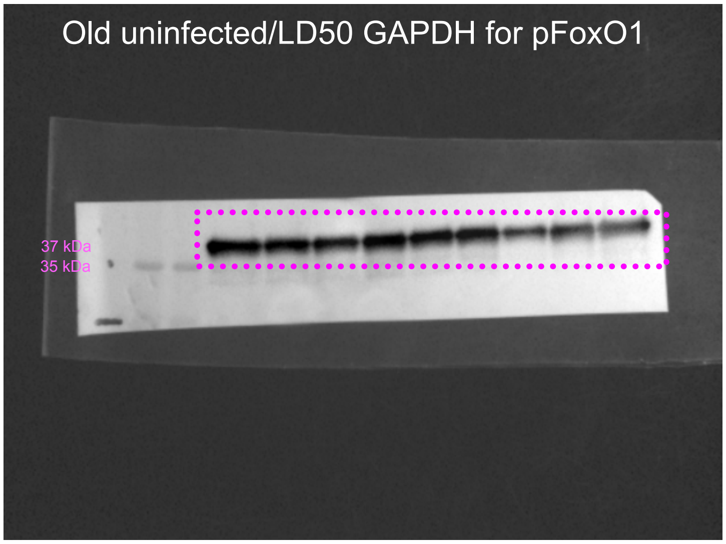

### Supplemental Figure 2

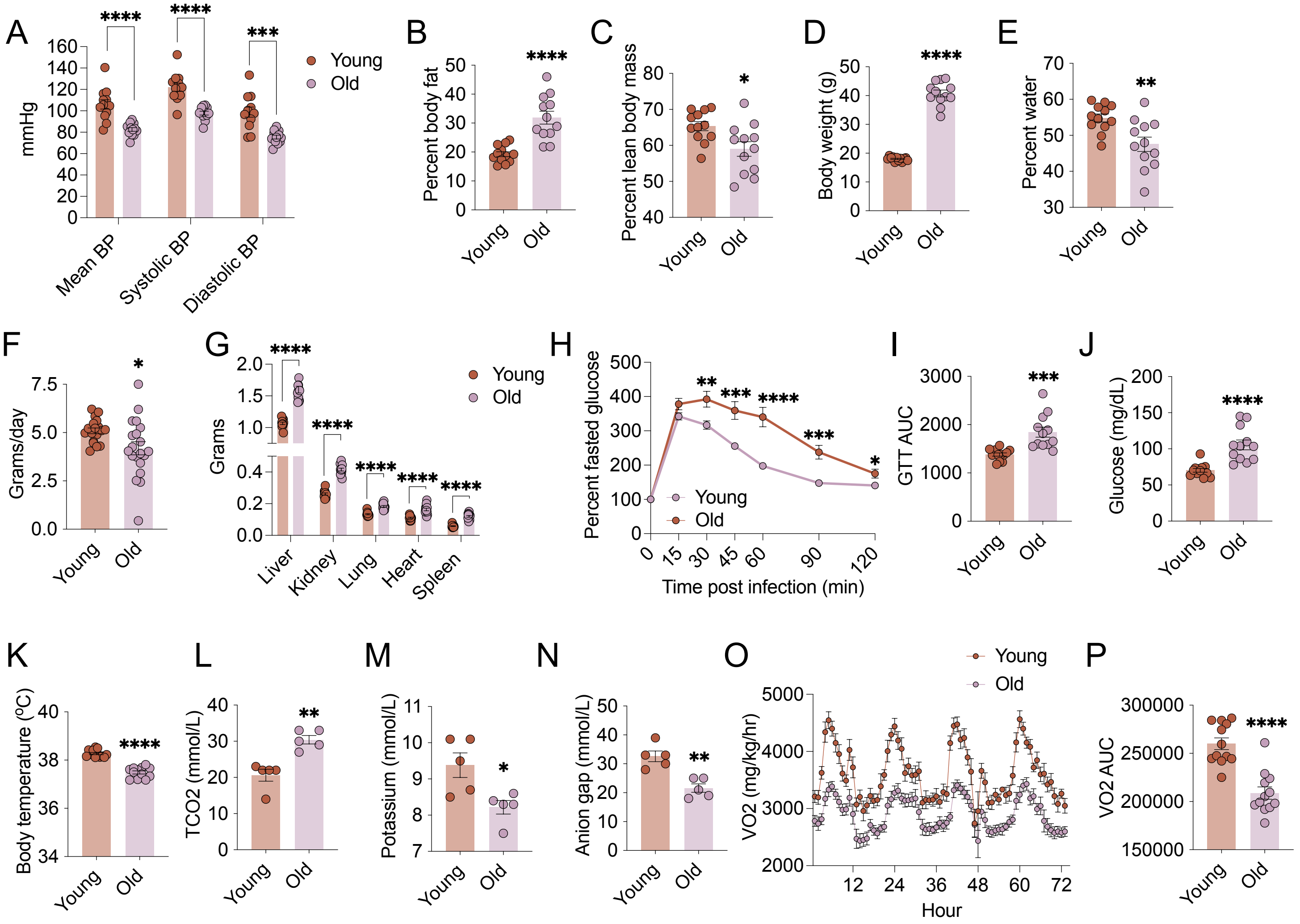

### Supplemental Figure 3

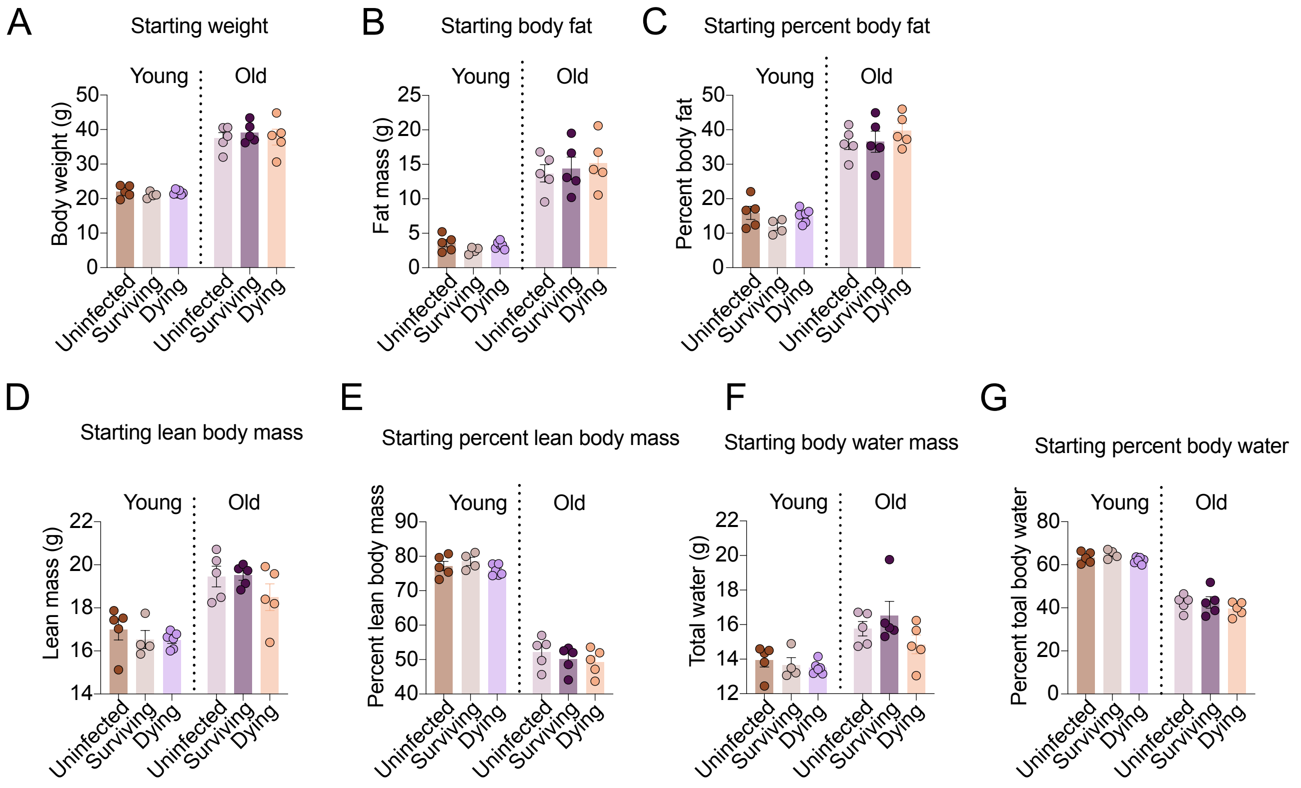

### Supplemental Figure 4

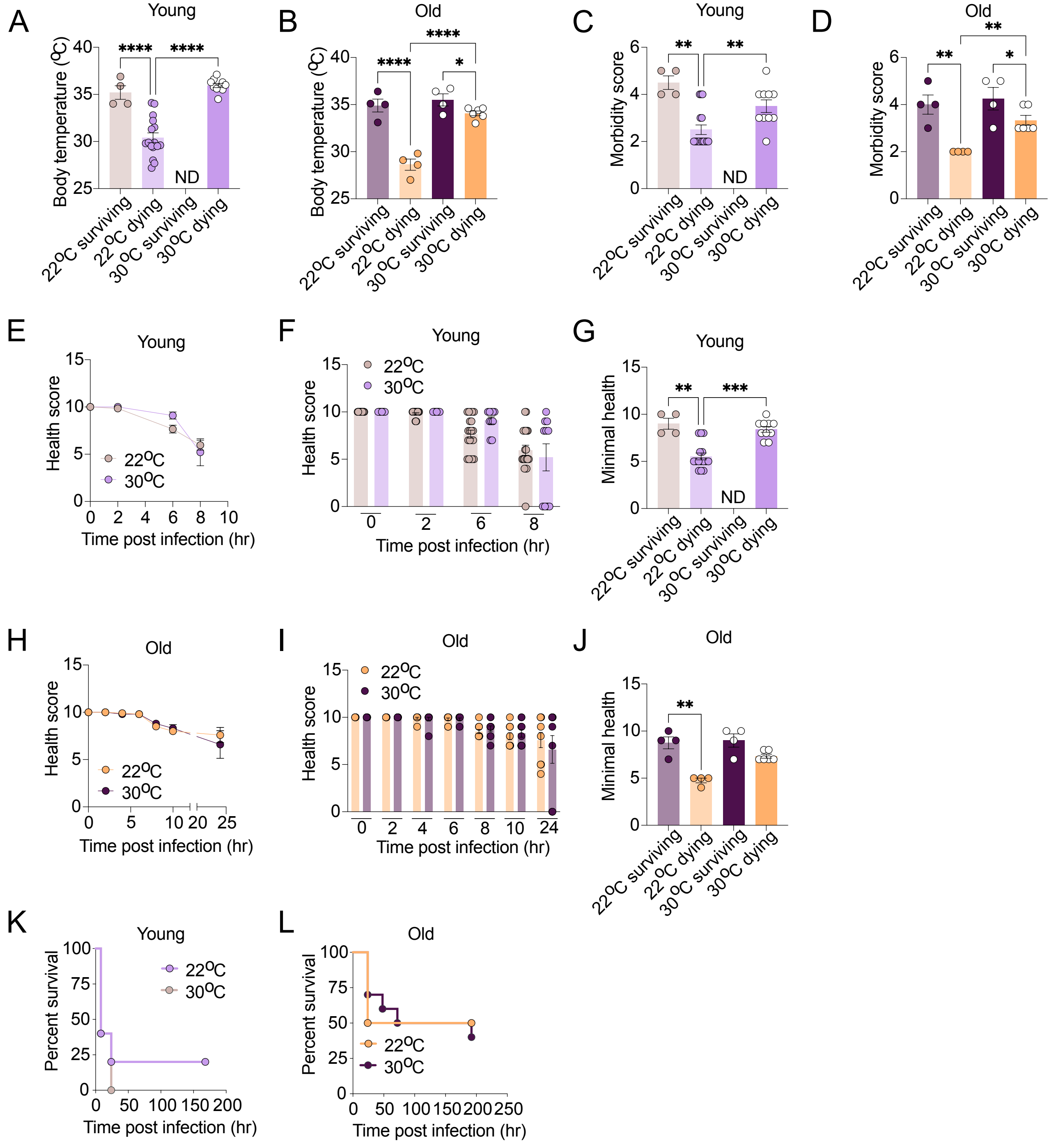

### Supplemental Figure 5

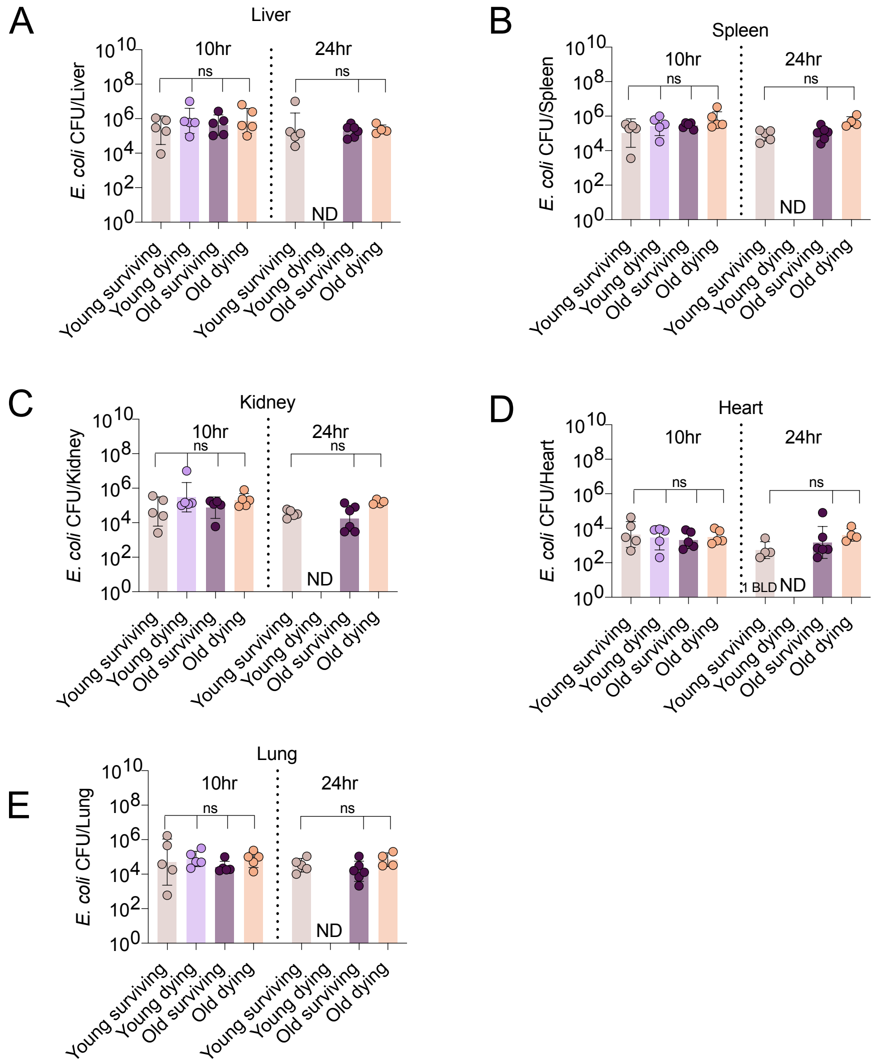

Supplemental Figure 6

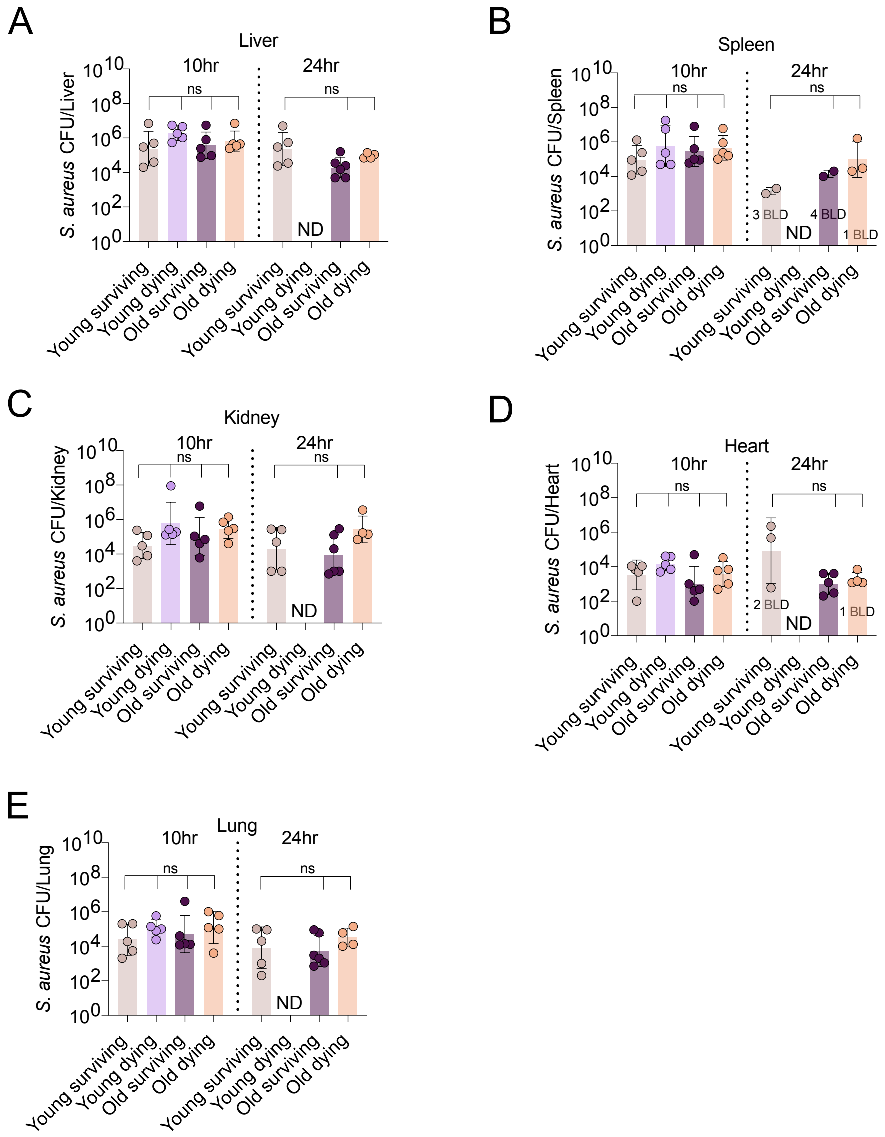

### Supplemental Figure 7

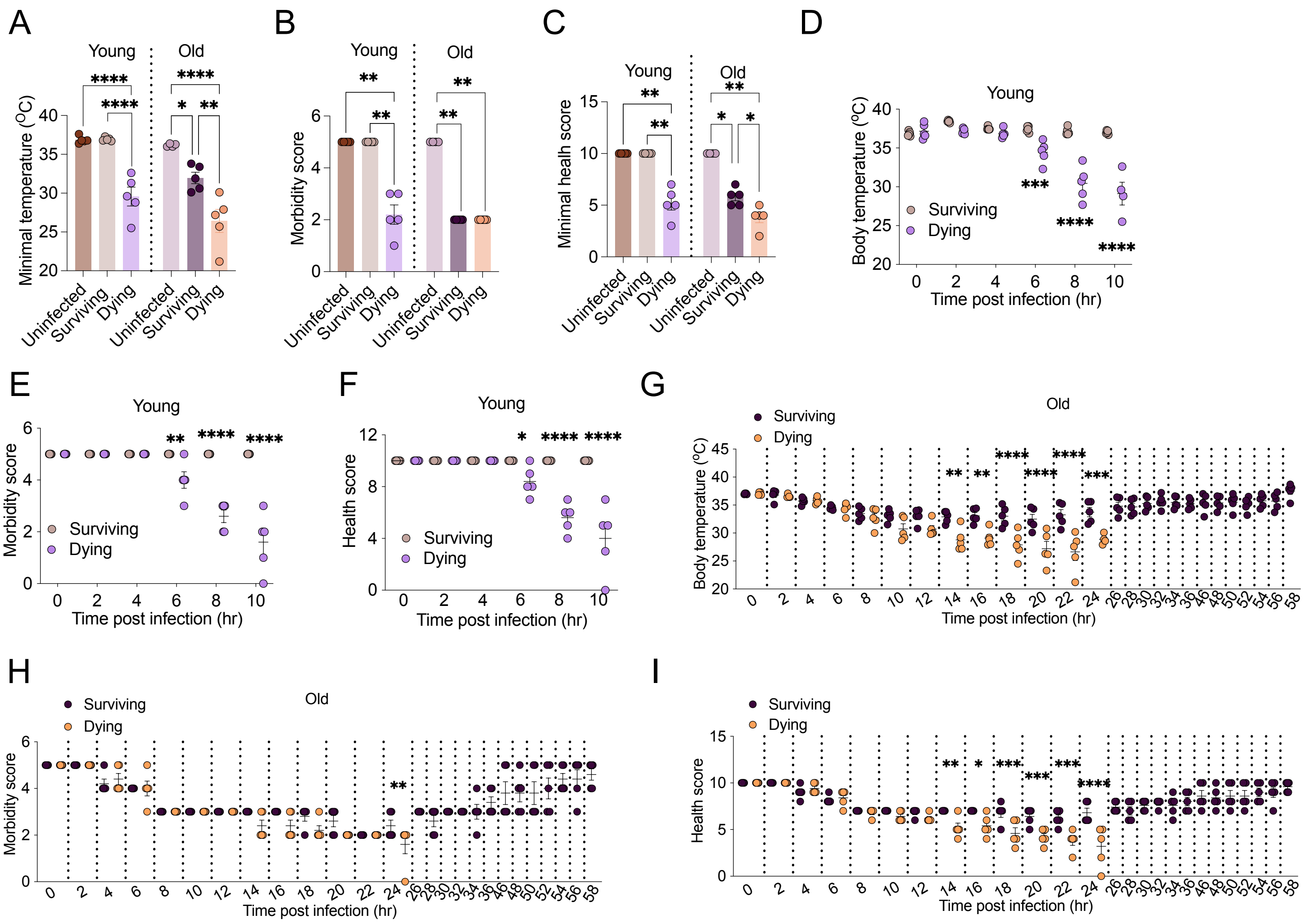

### Supplemental Figure 8

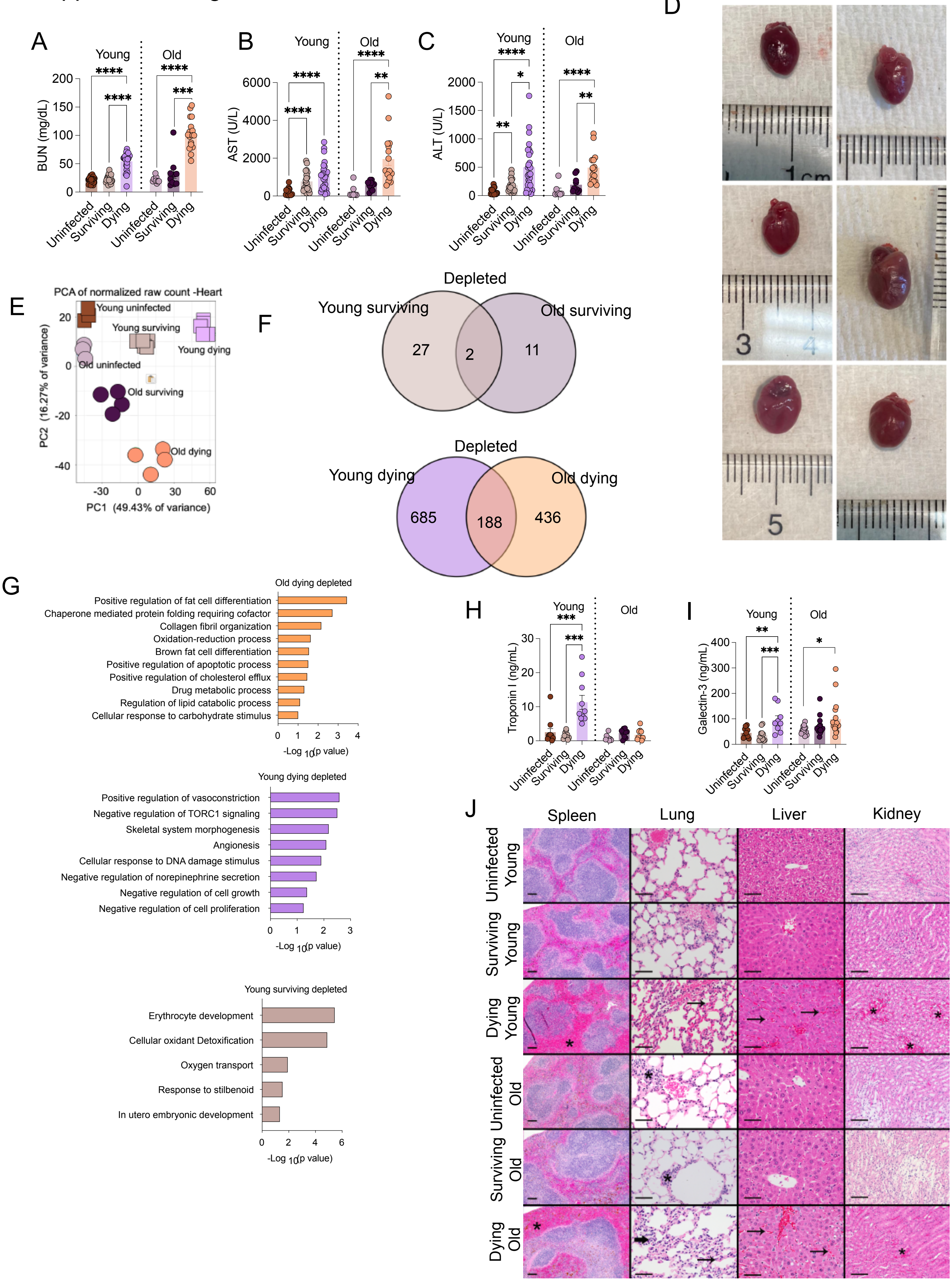

### Supplemental Figure 9

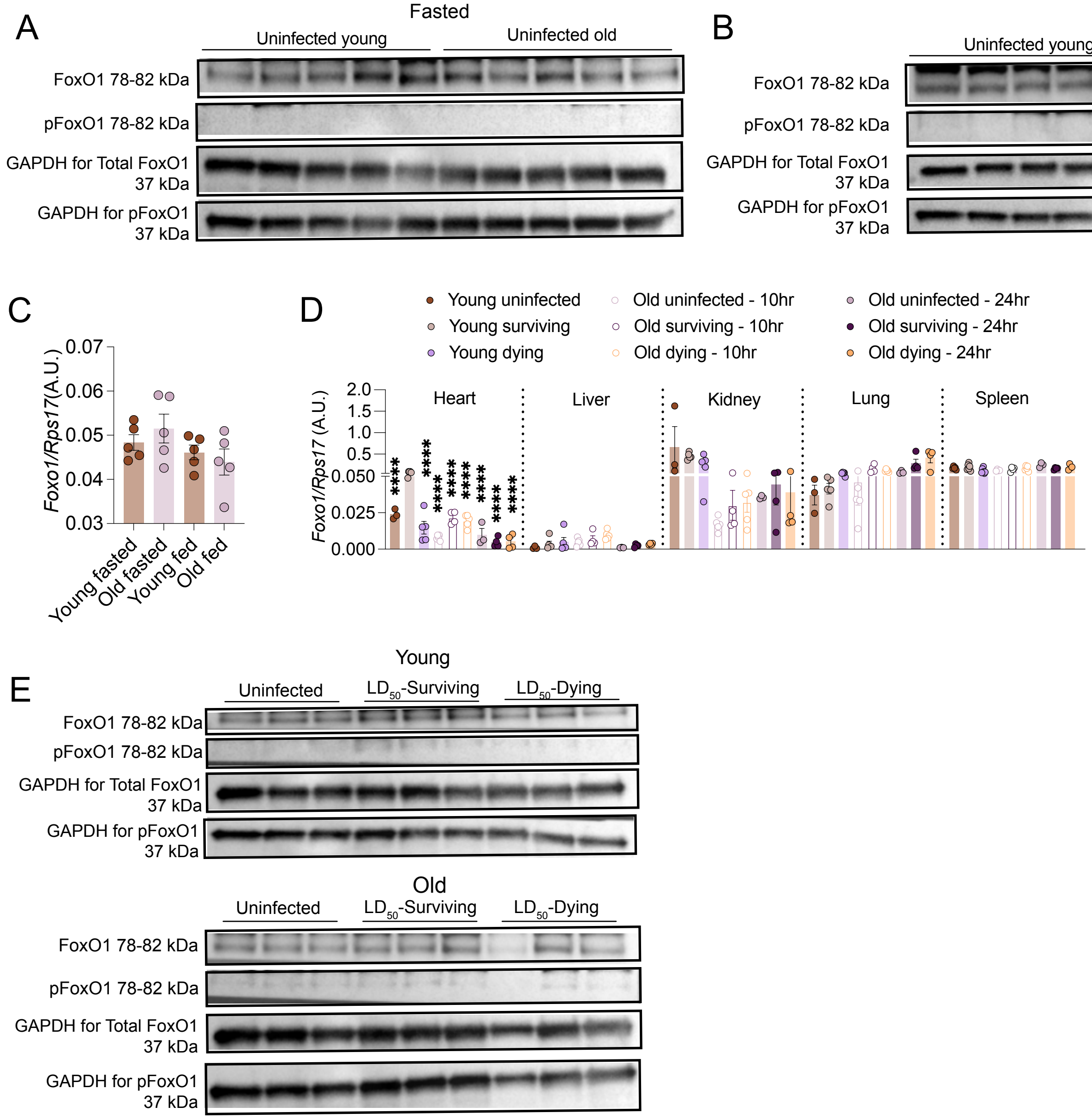

### Supplemental Figure 10

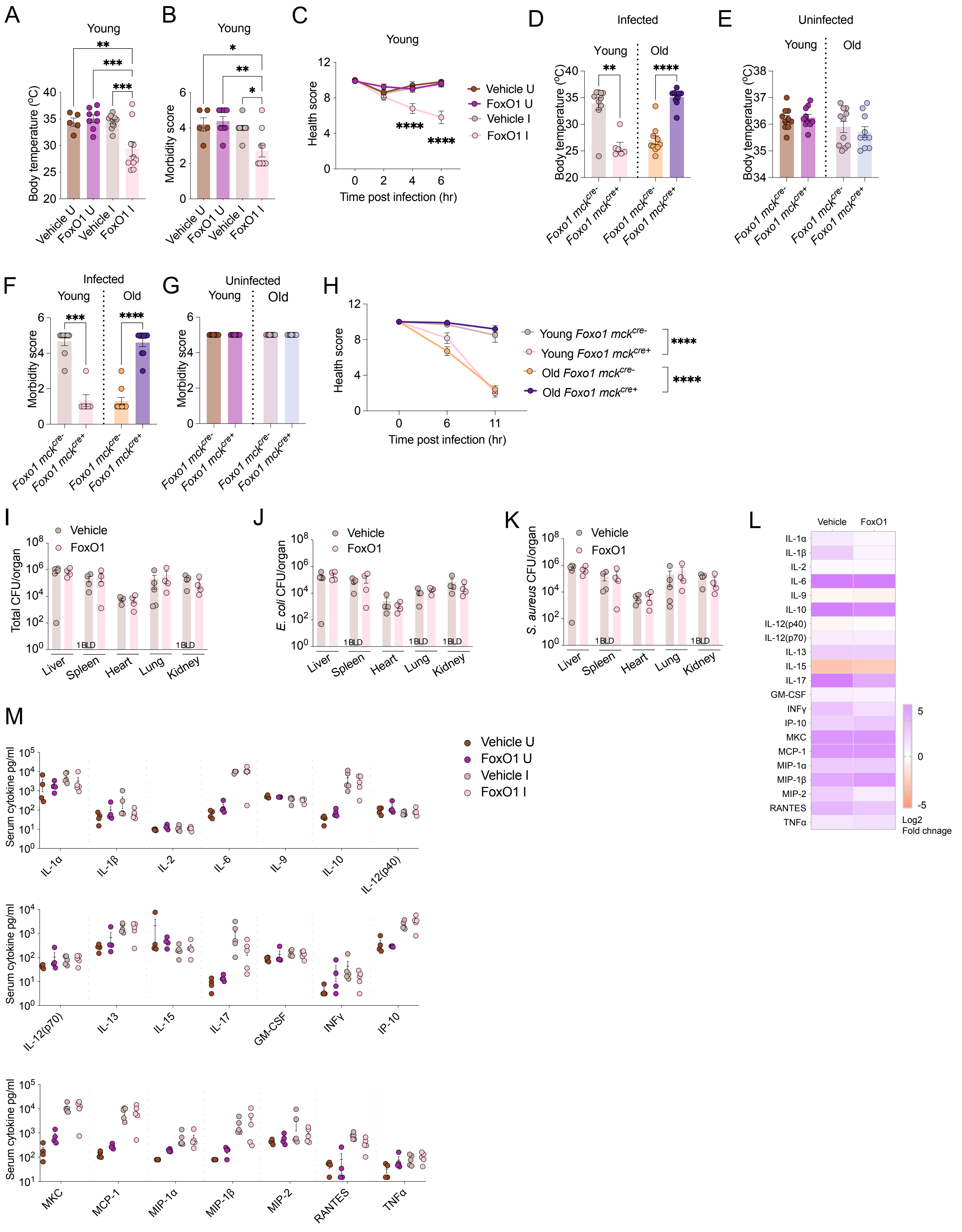

### Supplemental Figure 11

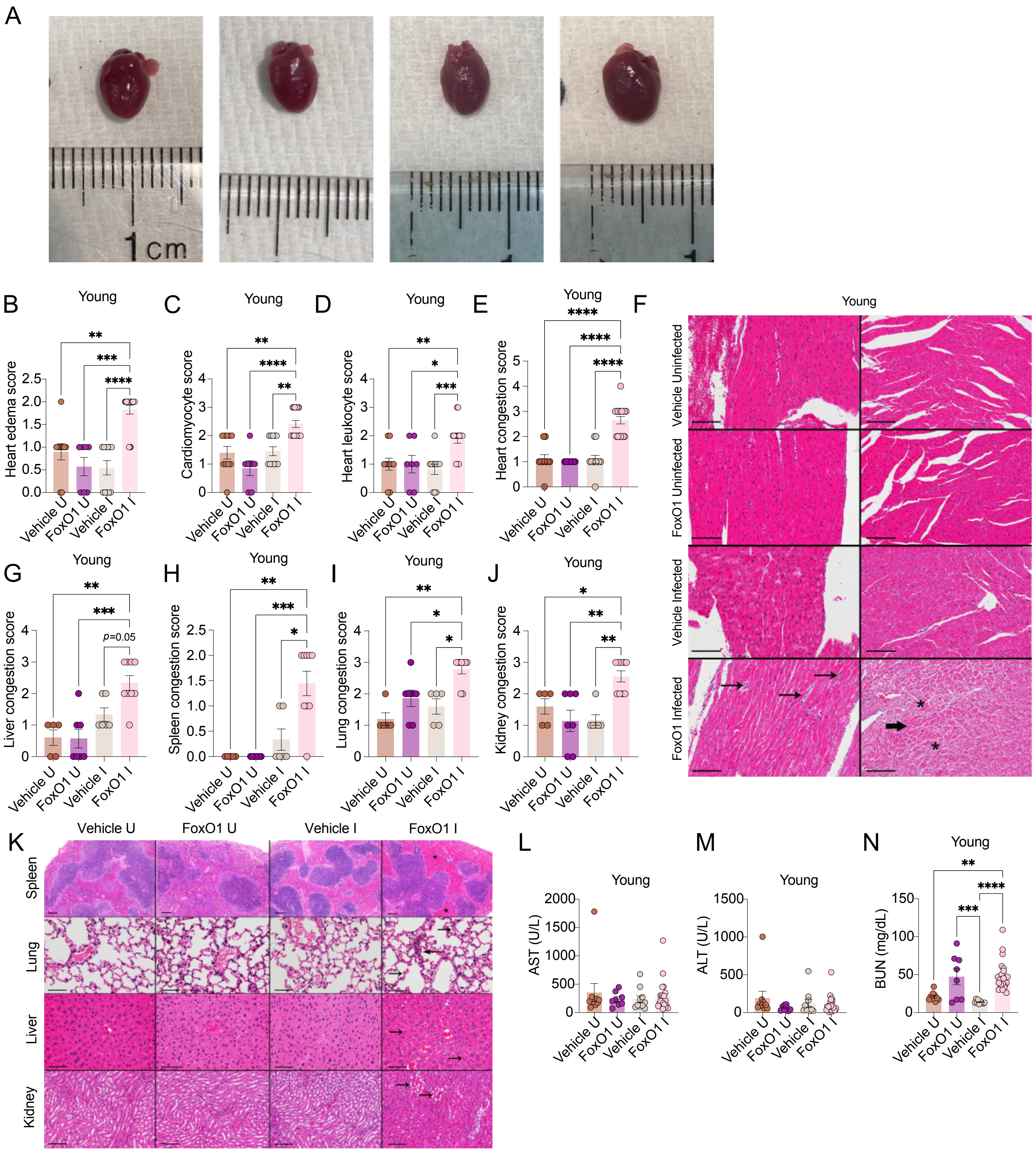

### Supplemental Figure 12

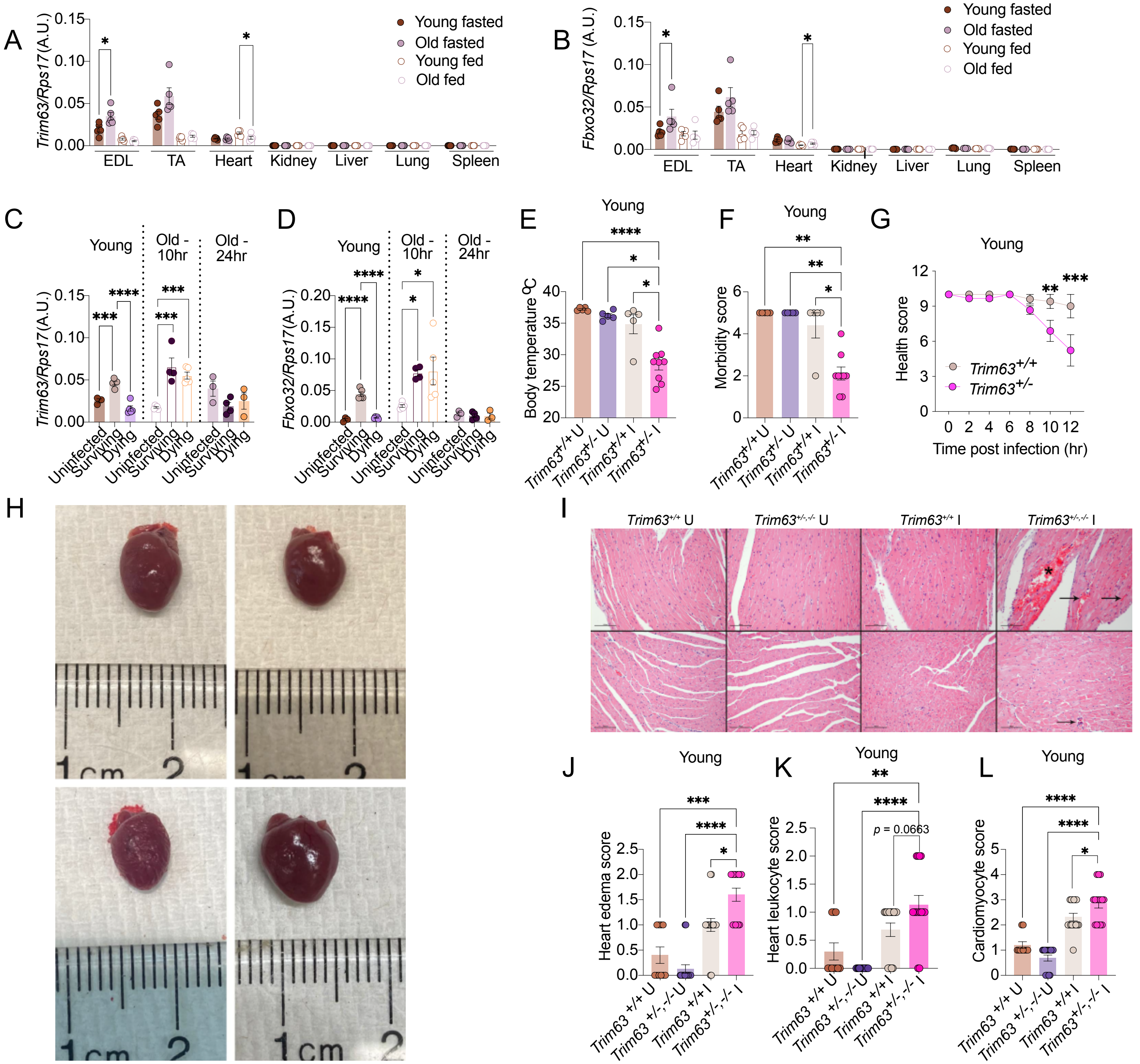

### Supplemental Figure 13

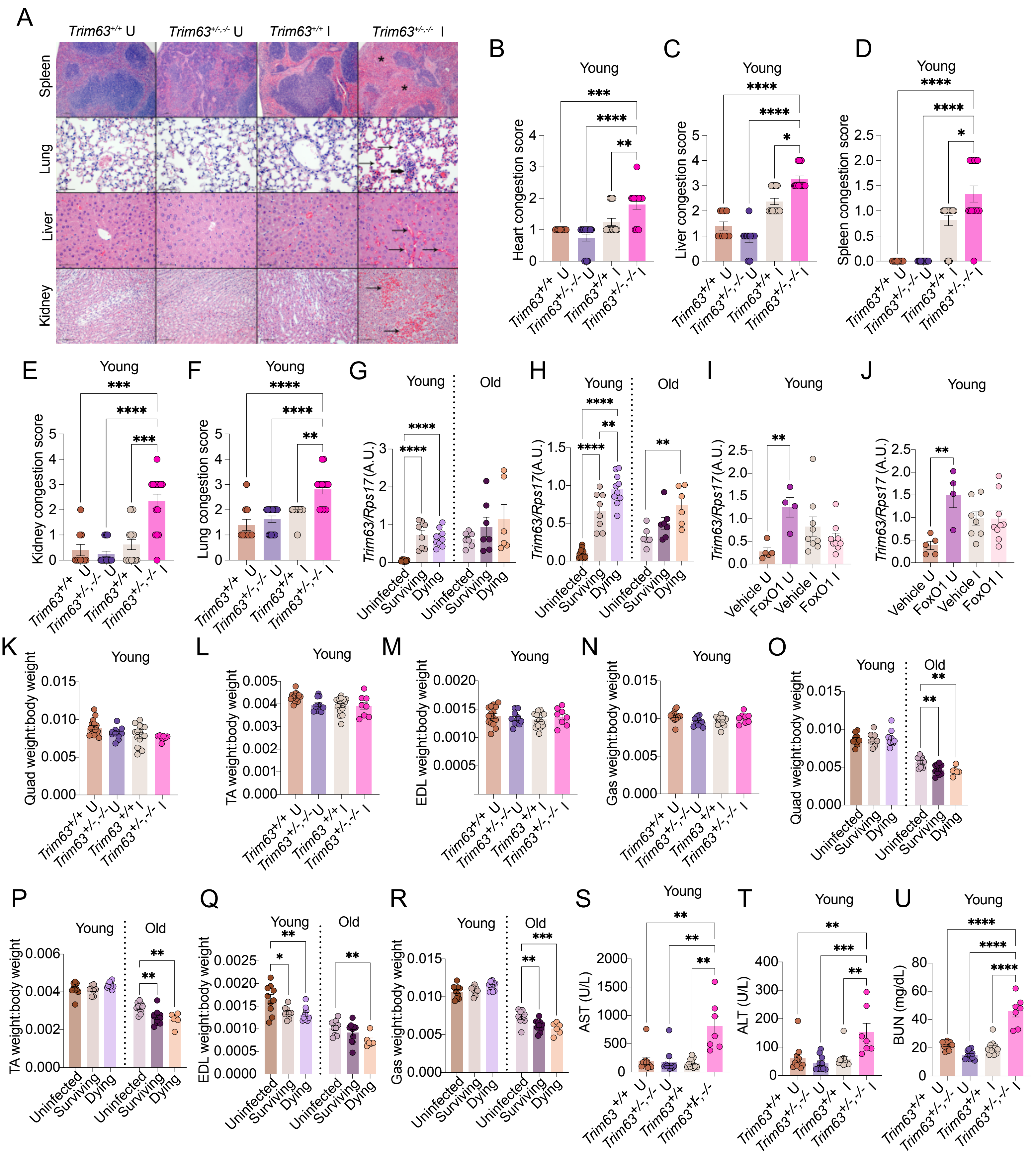

### Supplemental Figure 14

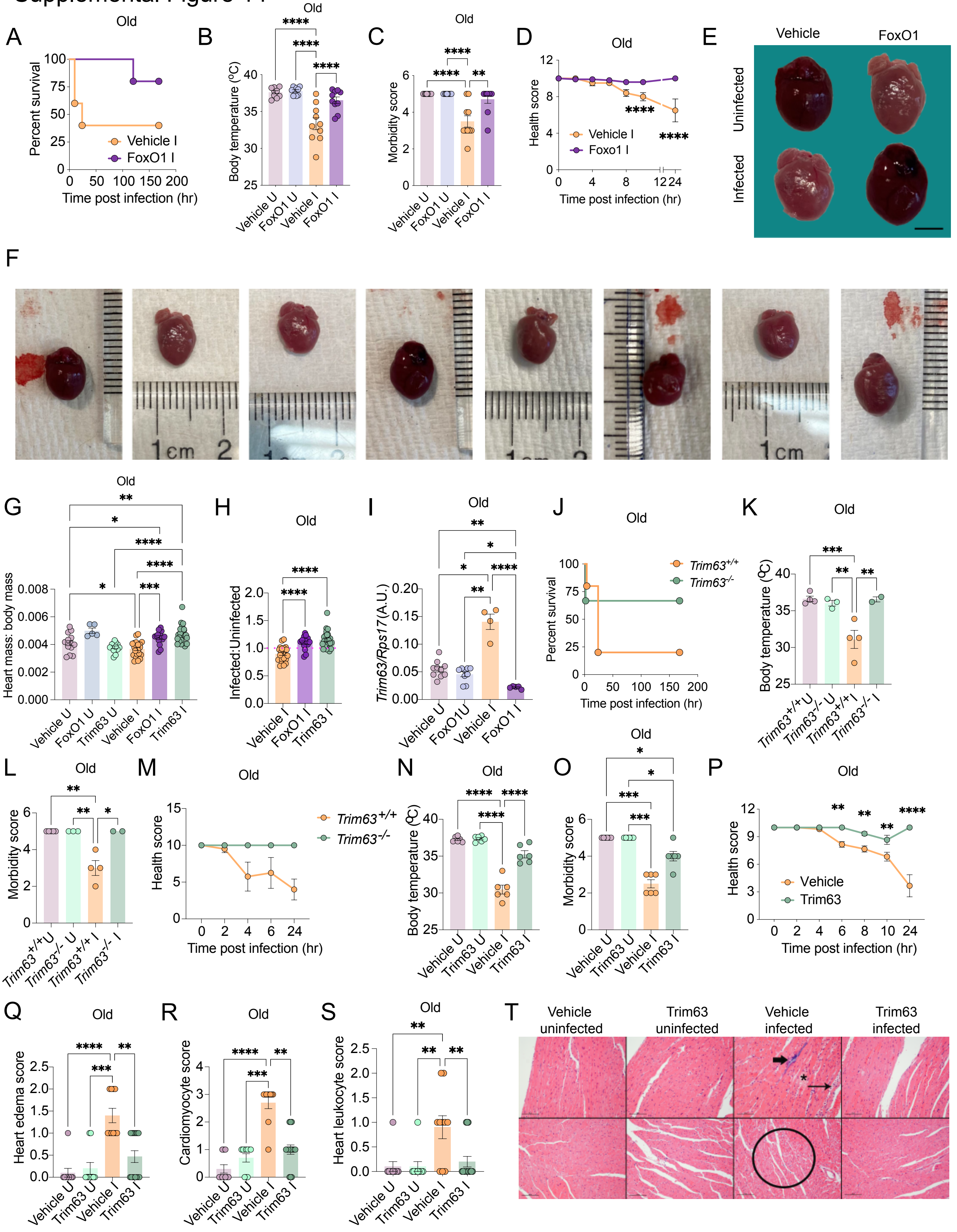

Supplemental Figure 15

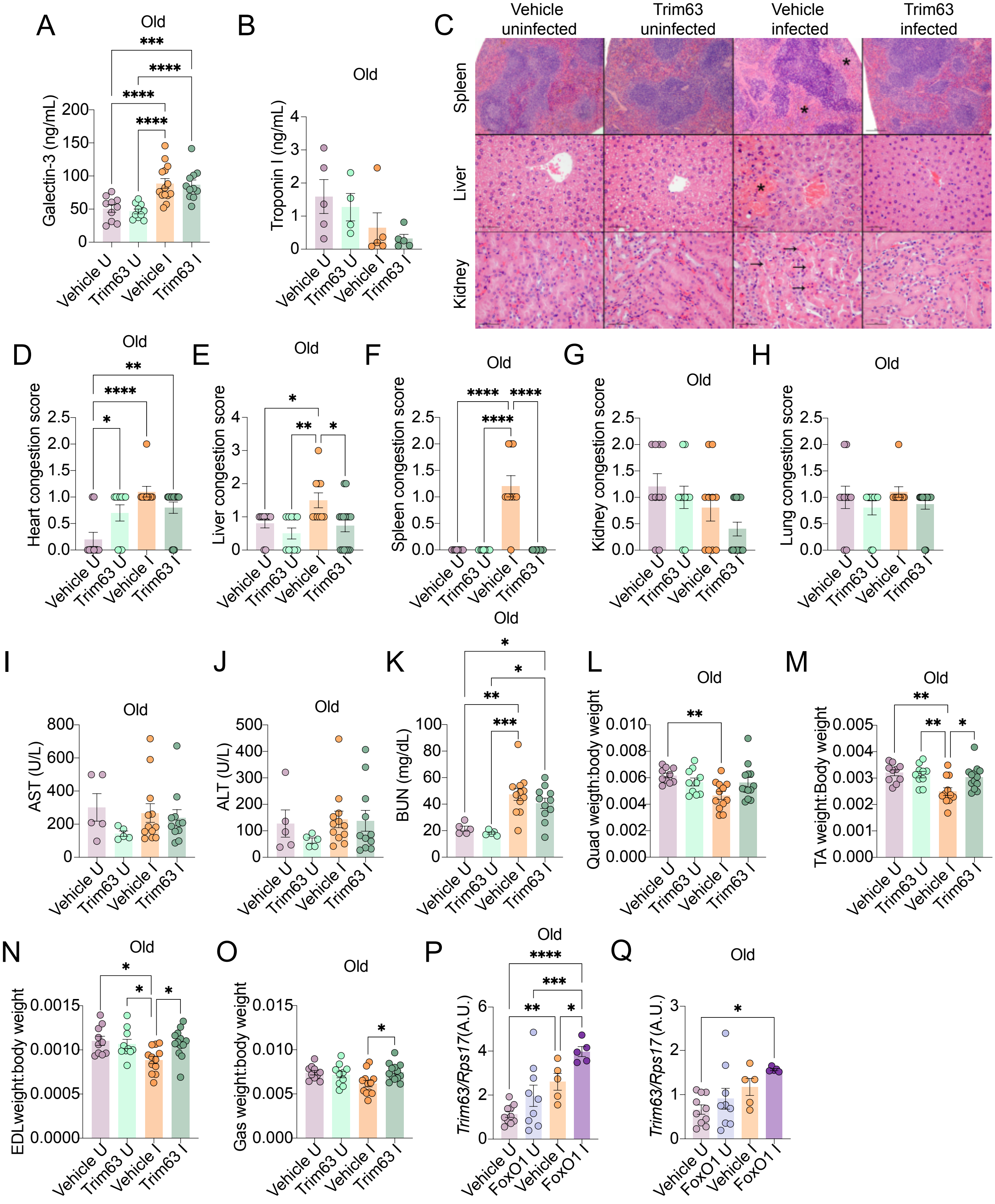
